# Structural Insights into Immature Dengue Virus-Like Particles Revealed by Cryo-EM and Molecular Dynamics Simulations

**DOI:** 10.1101/2025.01.21.631174

**Authors:** Venkata Raghuvamsi Palur, Guan-Wen Chen, Day-Yu Chao, Ya-Na Wu, Jedhan U. Galula, Chun-Hsiung Wang, Peter J. Bond, Jan K. Marzinek, Shang-Rung Wu

**Affiliations:** Department of Dentistry & Institute of Oral Medicine, National Cheng Kung University, Tainan, Taiwan; Bioinformatics Institute (BII), Agency for Science, Technology and Research (A*STAR), 30 Biopolis Street, #07-01 Matrix, Singapore 138671, Republic of Singapore; Graduate Institute of Microbiology and Public Health, College of Veterinary Medicine, National Chung Hsing University, Taichung, Taiwan; Department of Biological Sciences, National University of Singapore, Singapore, Republic of Singapore; Institute of Biological Chemistry, Academia Sinica, Taipei, Taiwan

**Keywords:** Dengue virus-like particles, cryo-EM, CHO-K1 cell line, molecular dynamics simulations, maturation transition pathway

## Abstract

The lack of efficacious vaccines against dengue (DENV) infections imposes an enormous burden on global health and the economy. Virus-like particles (VLPs), such as mature DENV VLPs (mDVLPs), have been shown to induce broadly neutralizing antibodies, making them promising next-generation vaccine candidates. However, the limited structural details have restricted efforts to engineer VLPs to attain the desired biophysical and immunological properties. In the current work, we present the cryo-electron microscopy (cryo-EM) structure of immature dengue serotype 2 VLP (imD2VLPs), revealing an architecture composed of a glycoprotein layer with prominent spikes in a T=1 arrangement. These spikes, composed of envelope (E) and precursor membrane (prM) protein heterodimers capped by pr domains, closely resemble immature flavivirus particles. Complementing the static structural details, we performed multiscale molecular dynamics (MD) simulations to elucidate the functional dynamics of imD2VLPs in the context of different lipid envelope compositions. Additionally, MD simulations uncovered the transition pathway between our previously solved mature VLP structure and immature VLP from this work. Here we show that VLP maturation involves a simple sliding-rotating motion without any clashes between E proteins. Our results also indicated that lipid composition plays a critical role in VLP stability, with phospholipid-dominant environments providing greater stability than diacylglycerol-rich vesicles. In addition, we demonstrated enhanced production efficiency of VLPs by generating a stable mammalian cell line using CHO-K1 cells. These findings enabled us not only to predict and manipulate the immunogenic properties of dengue VLPs but also underscored the potential of VLPs as a simplified and manageable model for investigating the structural basis of the dengue virus more effectively.

## INTRODUCTION

The global spread of dengue virus (DENV), a flavivirus primarily transmitted by *Aedes aegypti* and *Aedes albopictus* mosquitoes, leads to approximately 390 million infections annually^1^. DENV strains are classified into four distinct serotypes (DENV1 to 4)^2^. The clinical manifestations of DENV infection range from asymptomatic or mild febrile illness to severe conditions such as dengue hemorrhagic fever (DHF) and dengue shock syndrome (DSS), posing significant public health and economic burdens worldwide^3,4,5^. Since early 2023, the World Health Organization (WHO) has identified a resurgence of dengue cases and deaths in endemic areas, with the virus spreading to previously unaffected regions (WHO, 2024).

Given these increasing global health threats, developing an effective vaccine against dengue is becoming more urgent. However, this endeavor is challenging mainly because of the risk of “antibody-dependent enhancement” (ADE)^6^. ADE occurs when non-neutralizing antibodies from a prior infection help the virus to enter host cells, potentially exacerbating the disease. The currently recommended WHO vaccine has demonstrated suboptimal efficacy, particularly in DENV-naïve individuals, highlighting the urgent need for a safe and robust vaccine that provides balanced protection against all serotypes.

Virus-like particles (VLPs), which mimic viral antigenicity without containing the viral genome, represent a promising approach for vaccine development. They are advantageous due to their immunogenicity and potential for economical production across various systems. Several studies have shown the successful production of DENV VLPs in different expression systems, including plants^7,8^, insect cells^9,10^, bacteria^11^, *Pichia pastoris*^12–14^, silkworm transfected with recombinant baculovirus genome^15^ and mammalian cells^16–18^. Despite encouraging results, while many of these vaccine candidates have induced neutralizing antibodies and/or protection in animal models, including non-human primates^19^, detailed morphological studies using electron microscopy have not been fully elucidated.

Our previous studies^20^ have demonstrated that VLPs derived from dengue serotype 2 (mD2VLPs) can generate higher and broader neutralization antibodies against all four serotypes in mice experiments, demonstrating that these VLPs have the potential to be vaccine candidates for preventing DENV-induced diseases, as confirmed by cryo-electron microscopy (cryo-EM). Notably, the absence of pr peptide in mD2VLPs reduced the ADE effect, thus increasing their suitability as safe vaccine candidates. A crucial gap in designing optimal vaccine antigens lies in understanding the immature state of dengue VLPs and the functional significance of noncleaved prM proteins. Therefore, it is essential to investigate the potential impact of prM within the VLP system, compare the structure of VLPs at different stages to those of native virions, and consider their transient nature during maturation within the VLP framework.

DENV maturation is facilitated by furin, a protease primarily located in the trans-Golgi network (TGN). During this process, genomic RNA bound to C proteins forms a nucleocapsid core within the endoplasmic reticulum (ER), which acquires an envelope featuring 60 icosahedrally arranged trimeric spikes of pre-membrane (prM) and envelope (E) heterodimers, leading to the formation of an immature virus measuring 60 nm in diameter^21^ ^22^ ^23^. As the virus transits the acidic compartments of the secretory pathway, significant conformational changes enable furin to cleave prM into pr and M. This cleavage transforms the spikes from 60 trimeric prM/E heterodimers into a smooth structure of 90 heterodimers with an exposed furin site.

After furin cleavage, the pr remains bound to the E protein, effectively preventing premature membrane fusion until it is shed during the mature virus’s release through exocytosis. The mature virion, measuring 50 nm in diameter, contains 180 copies of E and membrane (M) protein heterodimers, which are presented as 90 head-to-tail homodimers. Three of these homodimers are positioned parallel to each other, forming a raft structure^21^ ^24^, which is ready for membrane fusion at low pH within the endosome.

Despite advances in cryo-EM have elucidated the significant structural transitions from immature particles with 60 prominent spikes to the mature virion’s smooth surface^25^, the transient nature of the dynamics remains a crucial aspect to study. Molecular dynamics simulations (MD) are useful in this regard, as they enable the study of a system’s motions at the atomistic (all-atom simulations) or residue (coarse-grained simulations) level^26^, providing information that may be unavailable from state-of-the-art experimental techniques alone.

The current study furthers our understanding by elucidating the structure and organization of the immature VLP, which undergoes global structural changes akin to the infectious virion. The cryo-EM structure of imD2VLPs produced by CHO-K1 mammalian cell line revealed a distinctive spiky surface, reflecting their immature state characterized by 20 trimeric prM-E protein protomers, compared to 60 in the case of virion. Additionally, multiscale MD simulations revealed the complex functional dynamics and the structural changes that characterize the likely VLP maturation pathway. The integration of cryo-EM with intricate computational modeling provides novel insights into the maturation processes of VLPs, which hold significant promise for the development of VLP-based vaccines.

## MATERIALS AND METHODS

### Plasmid and cells

The design of plasmid has been described previously^20,27^. Specially, the uncleaved-prM plasmid was generated by mutating the furin cleavage site of prM in pVD2i plasmid, which was deigned to comprise a gene coding for prM protein, 80% and 20% of C-terminus of E proteins from DENV-2 (Asian one genotype, strain 16681) and Japanese encephalitis virus (strain SA14-14-2), respectively. The plasmid was then transfected into mammalian cell lines CHO (Chinese hamster ovary)-K1-2-6-7 cells by using lipopolyamine (Transfectam; Biosepra, Villeneuve-la-Garenne, France) according to the instructions supplied by the company. The cells were routinely cultivated in suspension in DMEM medium supplemented with 10% Fetal Bovine Sera (FBS), 1% antibiotic-antimycotic, 1% Minimum Essential Medium (MEM) Non-Essential Amino Acids (NEAA) and 1% sodium pyruvate in a humidified atmosphere of 5% (v/v) CO_2_ at 37 °C. Cell cultures were maintained by seeding the cells at a low density.

### Harvesting and purification of imD2VLPs

For harvesting imD2VLPs, the CHO-K1-2-6-7 cells were seeded 5.0 x 10^6^ cells into T150 flask and cultured overnight. After the cell adherence, then medium was changed to serum–free maintenance medium (pH 7.4) containing 2% GlutaMAX^TM^-I, 1% sodium pyruvate, 1% MEM NEAA, 1% antibiotic-antimycotic and 1% cholesterol followed by transferring T150 flask to 28°C for additional 6 days. The culture supernatant harvested from cell line were clarified by centrifuging with 10,000 rpm at 4°C for 30 mins. The clarified superannuant was further applied for 20% sucrose cushion purification at 4°C at 25,000 rpm for 16 hrs. The purified VLPs pellets were resuspended in TNE buffer (50 mM Tris-HCl, 100 mM NaCl, 1 mM EDTA) and recovered at 4°C overnight. The recovered VLPs solutions was applied to a 5 to 25% (wt/wt) sucrose gradient (prepared with TNE buffer) and then centrifuged at 4°C at 25,000 rpm for 3 hrs in the Beckman SW-41 rotor. The fractions were collected and the level of E antigen in each fraction was determined by ELISA (see below). The pooled imD2VLPs containing fractions were concentrated and replaced with TNE buffer, using Amicon® Ultra 0.5ml centrifugal filters (100K MWCO) (Merck Millipore, Germany).

### Antigen capture ELISA

A conventional antigen-capture Enzyme-linked immunosorbent assay (ELISA) for quantifying imD2VLPs from the harvested culture supernatant was performed essentially as previously described^20^. Briefly, rabbit anti-DENV-2 VLP serum in coating buffer (28.3 mM Na_2_CO_3_ and 71.4 mM NaHCO_3_) was coated on 96- well plates, and blocked with 1% FBS in Phosphate-Buffered Saline (PBS) for 1 hr at 37°C. Subsequently, the VLP antigen in blocking buffer was added to wells. The bound antigens were detected by using anti-DENV2 MHIAF at 1:5000 followed by Horseradish peroxidase-conjugated goat anti-mouse IgG (Leadgene, Taiwan) at 1:6000. After additional incubation 37°C for 1 hr, the reactions were developed with 3,3’5,5′-Tetramethylbenzidine (TMB) substrate. The reaction was stopped with 2 N H_2_SO_4_, and absorbance was measured at 450 nm using ELISA reader (MULTISKAN GO, Thermo Scientific, Waltham, MA).

### Cryo-EM and 3D reconstruction

The immature DENV VLPs (imD2VLPs) structure was solved using single particle cryo-EM. The procedure followed the standard procedure. In particular, the 4μl purified imD2VLPs were loaded to glow-discharged holey carbon grid (Quatifoil GmbH, Germany). After 1 min in a high humid environment, the *g*rid was blotted by filter paper to remove the excess solution and then flash-frozen in liquid ethane bathed by liquid nitrogen implemented by the Vitrobot station (Mark IV, FEI). The frozen grid was stayed in liquid nitrogen to keep in low temperature and then transferred onto the precooling cryo-transfer holder. The micrographs images were captured at a magnification of 73,000X, resulting in a pixel size of 1.3785 Å per pixel by FEI Talos Arctica transmission electron microscope (ThermoFisher) at an accelerating of 200kV. During capturing of the images, each image produces 50 frames per 2.5 seconds, and these 50 frames aligned using cryoSPARC software^28^ to increase the signal of the image. Accordingly, the micrographs were patch-CTF corrected, the 2D classification was performed to reveal the heterogeneity and to remove junk particles. In the first run of 2D classification, 45 particles were manually picked, and the most 2 populated classes were selected for template-based particle picking. There were 12,563 particles which were auto picked from 350 micrographs. In the second run of 2D classification, 3,512 single particles with non-spherical features were discarded, resulting in 9,051 particles for creating a starting model *de novo* and the following homogeneous refinement (**Supplementary Fig. 1**). Resolution of the reconstruction was estimated from a Fourier shell correlation curve, using the gold standard method.

### Plastic embedded section specimen

The pelleted cells were embedded in resin-ethanol mixture with a gradually increasing ratio of Lowicryl to ethanol and in pure Lowicryl for the final infiltration. The plastic block was trimmed for ultrathin sectioning to 90 nm in thickness using diamond knife. Individual section was picked up on a carbon coated formvar copper slot grid (EMS, catalog number FCF2010-Cu). The grid was stained with uranyl acetate/lead citrate and imaged with JEM-1400 microscope.

### Coarse-grained model of imD2VLP

The initial of imD2VLP model was constructed by fitting the trimeric E-prM protein complex from the immature DENV serotype 2 virion structure (PDB: 4B03)^25^ into the imD2VLP cryo-EM density map. Full-length complete atomistic models of E and prM proteins were constructed with sequences outlined in the above experimental section. Modeller 9.10 was used to construct the homology models of the E-prM protein complex using immature DENV virion structure from PDB (PDB:4B03)^29^. A single unit of the E-prM heterodimer structure was superimposed onto the 60 E-prM protein units of the initial full immature VLP model composed backbone atoms. The resulting atomistic immature VLP model was converted to a Martini 3.0 coarse-grained (CG) model with an elastic network (EN) to retain higher order protein structure^30^. We applied the EN to protein backbone beads using a distance of 0.5 to 0.9 nm and a force constant of 1,000 kJ mol^-1^ nm^-2^. The EN was introduced only within each E protein domain (DI, DII and DIII). In the case of the prM protein, the EN was used only on the structured portion of the protein (residues 1 to 80), while TM regions of both E and prM protein had no EN as described previously^31^. We generated two lipid vesicles of radius 90 Å in agreement with the cryo-EM map (i.e., distance between center of lipid vesicle to the edge of inner leaflet) using the CHARMM-GUI Martini vesicle builder^32^ consisting of:

1. A phospholipid (PL) dominant composition, containing POPC (phosphatidylcholine 16:0/18:1) : POPE (phosphatidylethanolamine 16:0/18:1) : POPS (Phosphatidylserine 16:0/18:1) in a 60:30:10 ratio^33,34^.
2. A diacylglycerol dominant composition, containing DG (diacylglycerol 18:0/20:4) : SAPC (phosphatidylcholine 18:0/20:4): FA (fatty acid 18:1) in a 56:26:8 ratio in accordance with lipidomics data (reference to be added, manuscript under review).

In addition, for each lipid composition, two distinct vesicles with different total numbers of lipids were constructed. This was achieved by removing lipid molecules within either 1 Å (VLP_PL_-1 and VLP_DG_-1) or 1.5 Å (VLP_PL_-2 and VLP_DG_-2) from the protein CG beads. In total, four imD2VLP CG models were constructed. Each model was centered in a cubic box and MARTINI water was added together with 100 mM of NaCl on top of neutralizing the overall system charge^35^ (**Supplementary Table 1**). Each system energy was minimized for of 50,000 steps using the steepest descent algorithm. Subsequently, the system was subjected to 500 ns equilibrium simulation in the NPT ensemble with position restraints applied to protein backbone beads with a force constant of 1000 kJ mol^-1^ nm^-2^. Bond lengths between backbone beads were constrained using the LINCS algorithm^36^. Equations of motion were integrated using leapfrog algorithm with a 10-fs time step. The Velocity rescale thermostat^37^ and Berendsen barostat^38^ were used to maintain temperature and pressure at 310 K and 1 bar, respectively. Three independent NPT ensemble unrestrained replicas were run for 2,000 ns to 2,500 ns each (**Supplementary Table 1**). All CG simulations were run using Gromacs 2020.2 on 8 nodes containing 24 CPUs and 1 GPU each on the National Supercomputing Center (NSCC), Singapore.

### Targeted molecular dynamics (TMD) simulations

TMD simulations were carried out using Gromacs 2018.2^39^ with Plumed 2.6.4^40,41^ by applying a harmonic potential to the root mean square deviation (RMSD) with respect to the target structure of protein backbone beads^42,43^. Four independent force constants were used: 100, 500, 1,000 and 10,000 kcal mol^-1^ Å^-2^ of the additional RMSD harmonic potential bias with respect to the target structure (**Supplementary Table 2**). The RMSD harmonic potential was applied to E protein residues 1 to 495. In the case of prM protein, steering forces were applied to the TM region alone (residues 150 to 166). Every 1,000 steps of TMD simulation, the RMSD between the current and target structure was calculated. The transition was linearly triggered over the simulation time using the RMSD difference between the immature and mature VLP state totaling an RMSD of ∼58.7 Å. TMD was employed for 20,000,000 steps with a time step of 5 fs resulting in a 100 ns trajectory for each system. All TMD simulations were carried out for the VLP-1 system with the PL dominant lipid vesicle (**Supplementary Table 1**) and the same simulation settings were used as in the previous section.

### All-atom molecular dynamics simulations of glycosylated trimer embedded in lipid bilayer

The coordinates for a single trimeric E-prM protein of the immature DENV (PDB: 4B03) were used. Modeler 9.10^29^ was used to generate a homology model with E and M protein sequences as used in experiments. The lowest discrete optimized energy (DOPE) scored model structure with >97% amino acids in allowed regions of Ramachandran plot was used and embedded in a lipid bilayer composed of diacylglycerol (DG), phosphatidylcholine (PC) and fatty acid (FA) at 56:26:8 ratio as reported in Palur et al (reference to be added, manuscript under review). Glycosylation sites N153 and N67 sites was modelled using the CHARMM-GUI glycan builder^44^ according to the glycan composition of DENV2 reported in previous studies^45^ (**Fig. 7Dii**). The default protonation states at neutral pH were assigned to the E-prM ionizable residue using the CHARMM36m forcefield^46^. The trimeric unit was embedded into a lipid bilayer using the CHARMM-GUI membrane builder and solvated in a TIP3P water box^47,48^ plus 150 mM of NaCl which also served to neutralize the overall charge of the system. The membrane embedded E-prM system was subjected to 5,000 steps of steepest descent minimization followed by two and four steps of NVT and NPT equilibration steps respectively. After each equilibration step the force constant on protein backbone and lipid headgroup atoms was gradually reduced from 4,000 to 0 kJ mol^−1^ and 2,000 to 0 kJ mol^−1^, respectively. A total of 1.5 ns of equilibration simulation using a 1 fs (NVT) and 2 fs (NPT) time step was run using the leap-frog algorithm. During equilibration, a 310 K temperature and 1 bar pressure were controlled using the Berendsen thermostat^38^ and semi-isotropic barostat. The Particle mesh Ewald (PME) method^49,50^ was used for long range electrostatic interaction with a cut-off distance of 1.2 nm, and a switching function was applied after 1 nm. Production runs included Nosé-Hoover thermostat^51^ (303K) and semi-isotropic Parrinello-Rahman barostat^52^ (1 bar) with 1 ps and 5 ps coupling constants, respectively. In the production runs, the positions of the TM backbone atoms of E and M protein were restrained using a force constant of 1,000 kJ mol^−1^ nm^-2^. Restraints on the TM region were imposed to maintain the trimeric state of E-prM protein trimers akin to the immature state. Three independent production simulations of 100 ns each were carried out with restraints on the TM region alone. All simulations employed Gromacs 2020.2 on 4 nodes with 24 CPUs and 1 GPU on the National Supercomputing Center (NSCC), Singapore.

### Simulation analysis

To quantify the structural changes observed during TMD simulations driving the imD2VLP to mD2VLP transition, we measured three specific angles between either E protein monomers or dimers: ϴ, Φ and δ (**Supplementary Fig. 2**). We defined ϴ as the angle between vectors composed of E protein monomers as shown in Supplementary Fig. 2A, where each vector passes through center of mass (COM) of domain II and COM of domain III ectodomain of a respective E protein monomer. Φ was defined as the angle between two vectors, where the first vector is the line joining COM of DII and DIII of a E monomer and the second was a vector joining COM of the entire VLP and COM of DIII of the respective E protein monomer (**Supplementary Fig. 2B**). δ was defined as the angle between the vector connecting COM of DIII and DII of each monomer at time t=0 and the same vector at different time points during the TMD-1 simulation (**Supplementary Fig. 2C**). In all simulations, gromacs tools were used to perform principal component analysis (PCA), calculate RMSDs, contacts and cutoffs, and calculations of angles, while VMD, pymol and ChimeraX were used to visualize molecules and simulation trajectories.

## RESULTS

### Establishment of a stable cell clone continuously releasing prM -E containing VLPs

In our previous study, we investigated the production of mD2VLPs via transient transfection of COS-1 cells with plasmids encoding M and E proteins. This resulted in the induction of high titers of neutralizing antibodies against dengue virus following mD2VLP immunization in mice^20^. Our cryo-EM analysis of the secreted particles revealed that the mD2VLPs displayed T = 1 icosahedral symmetry, exposing highly overlapping cryptic neutralizing epitopes, thereby suggesting their potential as a vaccine candidate^20^. Building upon these results, our current work focuses on establishing a mammalian cell clone capable of producing dengue serotype 2 virus-like particles (imD2VLPs) more efficiently than traditional transient transfection approaches.

Inspired by the work of Konishi et al., who engineered a CHO-K1 cell line to produce high yields of Japanese encephalitis viral (JEV) antigens by modifying the prM/M cleavage site to reduce toxic fusion activity^53^, we pursued a similar strategy. Specifically, we transfected CHO-K1 cells with a cDNA encoding a chimeric E protein (80% DENV-2 and 20% JEV) alongside a mutated prM furin cleavage site to stably express DENV prM-E proteins. This modification inhibited furin cleavage and facilitated the assembly of immature dengue serotype 2 virus-like particles (imD2VLPs). Transmission electron microscopy (TEM) performed on thin sections of epon-embedded CHO-K1 cells at 72 hours post-trypsinization revealed the presence of spherical particles with electron-dense material closely associated with the plasma membrane (**Fig. 1A**). Antigen capture ELISA was used to quantify the release of E antigen from the cultured cells across multiple passages, demonstrating that the CHO-K1 cell clones stably secreted imD2VLPs for at least four passages (**Fig. 1B**).

**Figure 1.**
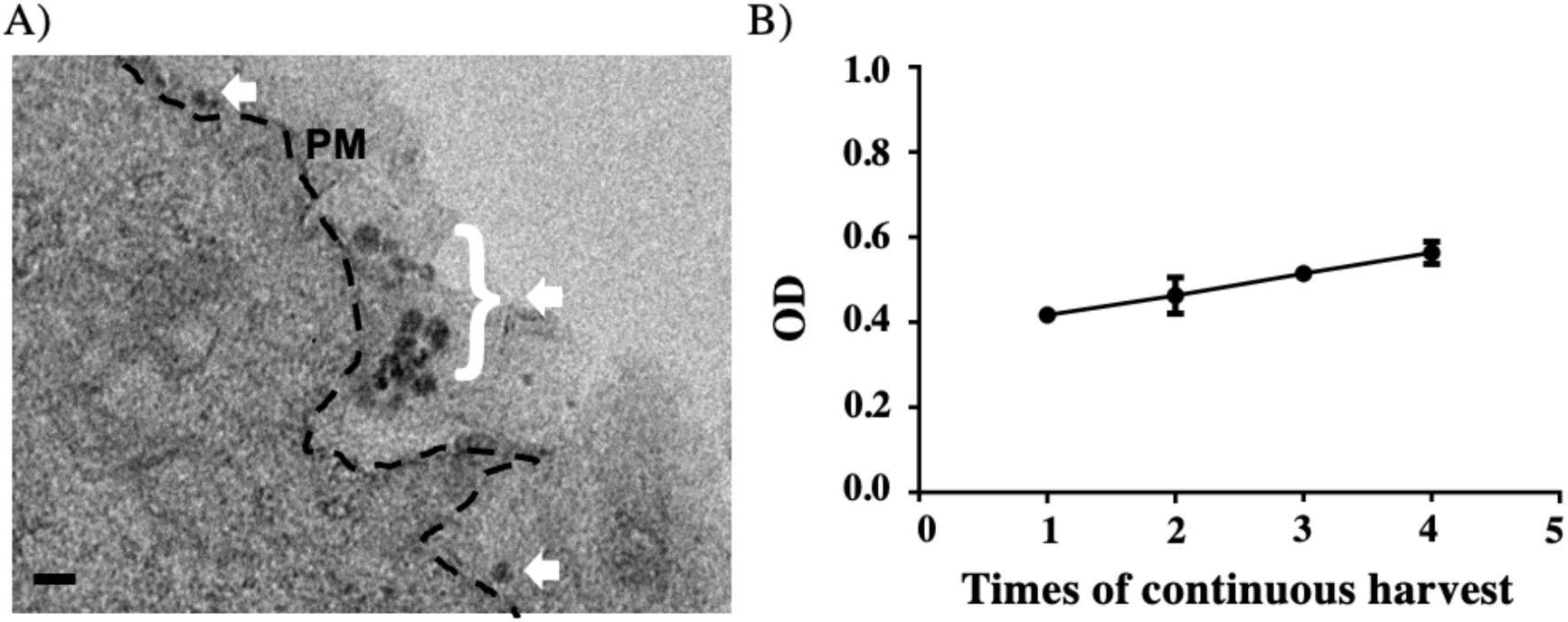
Quantification of immature dengue serotype 2 VLP (imD2VLPs) released from CHO-K1 Cells. **(A)** Transmission electron microscopy images of resin-embedded section of CHO-K1 cells, taken 72 hours post-infection (hpi), revealing spherical imD2VLPs (highlighted by white arrows). These electron-dense particles are prominently accumulated along the plasma membrane (PM), as marked by the dashed line. Scale bars=50 nm. (**B)** Quantification of imD2VLPs released from CHO-K1 cells. The culture medium containing the VLPs was harvested four times at six-days intervals. The culture medium of confluent cells was refreshed every six days, and no significant cell fusion or apoptosis was observed during the harvesting period. The quantity of imD2VLPs was measured via absorbance at 450 nm using an ELISA reader.

### Characterization of imD2VLPs

To further characterize the imD2VLPs, we purified them via sucrose gradient centrifugation and performed cryo-EM analysis. Cryo-EM imaging revealed that the concentrated imD2VLPs were monodispersed (**Fig. 2**), exhibiting a rougher surface compared to mature dengue VLPs^20^ appearing significantly smaller than natural immature flaviviruses^54,55^. Two-dimensional classification of imD2VLPs identified three distinct populations based on morphology and size (**Fig. 2; Supplementary Fig. 1**). Most of the population (76%) consisted of particles with a symmetrical, prominent protrusion surrounding the spherical lipid bilayer, while the second population (22%) exhibited surface irregularities and non-spherical lipid bilayers. A minority of the population (2%) comprised double-membrane vesicles ranging from 40 to 52 nm in size.

**Figure 2.**
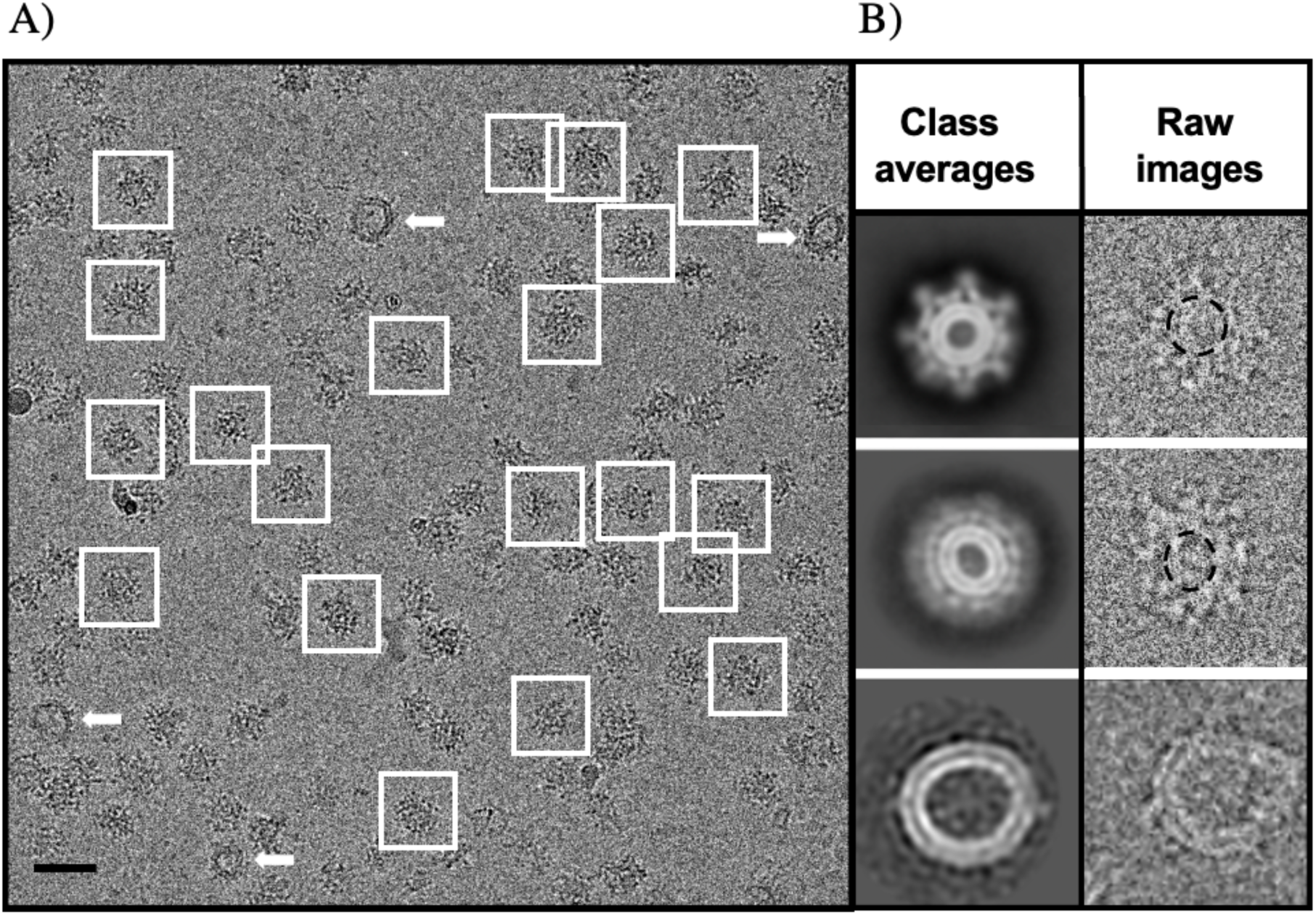
The cryo-EM image analyses of imD2VLPs. **(A)** Cryo-EM images of concentrated imD2VLPs show a monodispersed population of particles. Particles selected for further processing are highlighted with boxed regions. Double membrane vesicles are indicated by arrows. **(B)** Representative 2D class averages from reference-free classification display three distinct populations within the sample: the particles with symmetrically arranged protrusions that encapsulated the spherical lipid bilayer (top); the particles with less defined, blurry protrusions surrounding an imperfectly spherical lipid bilayer (middle); and the double membrane vesicles (bottom). The left panel shows the 2D class averages, while the right panel presents the corresponding raw images.

The cryo-EM reconstruction of the predominant population of imD2VLPs achieved an average resolution of 9.2 Å, revealing a multilayer architecture featuring 20 prominent spikes in a T = 1 arrangement (**Fig. 3**). The spikes measured approximately 41 nm in length, significantly larger than the ∼31 nm spikes observed in mD2VLPs^20^. Fitting the trimeric E-prM protein from the immature DENV serotype 2 virion structure (PDB: 4B03)^25^ into the imD2VLP cryo-EM map revealed that each spike consisted of three E-prM heterodimers capped by three pr domains. These surface protein interactions exhibited 3-fold symmetry, consistent with immature flavivirus structures, with notable features, including the buried furin cleavage site between E and pr domains (**Supplementary Movie 1**)^25,54–56^. The structure further highlighted steric hindrance at domain III (DIII) and amino acid 101 within the fusion loop, aligning with our previous observations that imD2VLPs elicit lower levels of fusion loop-specific and DIII conformation-independent antibodies compared to mD2VLPs^20^. These findings provide critical insights into the structural properties and unique features of imD2VLPs, shedding light on their potential use in vaccine development.

**Figure 3.**
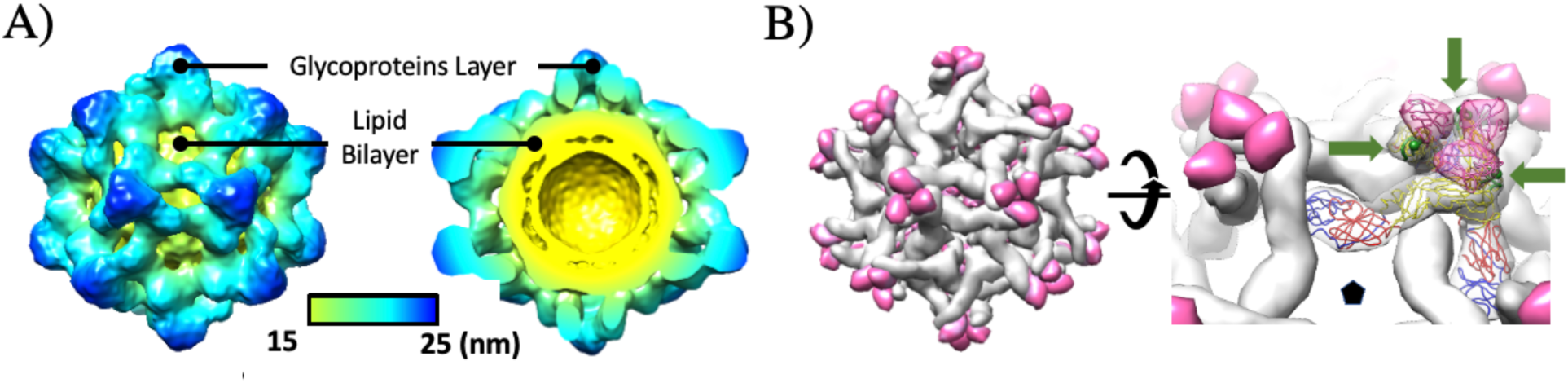
Cryo-EM structure of the imD2VLP. **(A)** The sharpened cryo-EM map of imD2VLP at 2.1σ, viewed from 2-fold axis (left), radially colored to indicate particle diameter. The cross-sectional view (right) reveals the multilayer architecture, consisting of the glycoprotein layer (blue-green) and the lipid bilayer (yellow). The color bar denotes the particle diameter. **(B)** Surface rendering (gray) of the fitted surface proteins (PDB: 4B03) reveals that the imD2VLP conforms to T=1 icosahedral symmetry. The spike is composed of a (prM/E)_3_ trimer, where the E protein exists as a heterodimer with prM. The cleaved pr peptide (magenta) behaves as a cap on the E protein (left). A closer examination of the fitted surface proteins discloses steric hindrance within domain III (DIII) of the E protein during antibody binding (right). The critical binding residue for DM25-3 interaction, amino acid 101 (shown by green spheres) in E protein, is positioned inside the spike, resulting in partial steric hindrance during antibody interaction (right). E protein monomers are color-coded by domain: domain I (DI) in red, domain II (DII) in yellow, and domain III (DIII) in blue. The pr peptides are shown in magenta, and the fusion loop (green) is indicated by green arrows. The pentamer denotes the 5-fold symmetry axis.

### CG Simulations reveal imD2VLPs intrinsic dynamics and lipid composition-specific morphological changes

To explore the intrinsic dynamics of imD2VLPs, we built several CG models of the entire imD2VLPs consisting of 20 trimeric protomers of E-prM protein. These were embedded in lipid vesicles of 90 Å radius according to the cryo-EM density map (**Fig. 4A-C**). We used two different experimentally determined lipid compositions with various numbers of lipids in different possible vesicle models to assess the stability of imD2VLPs and compared the results with cryo-EM micrographs (**Fig. 4B-C**). We studied PL dominant vesicles containing POPC:POPE:POPS in a 60:30:10 ratio^33,34^ (VLP_PL_) as well as DG dominant vesicles composed of DG:SAPC:FA in a 56:28:6 ratio (VLP_DG_) as revealed by the experiments using a COS-1 cell line (reference to be added, manuscript under review). We tested lipid vesicles with two distinct total numbers of lipids for both the PL and DG dominant lipid compositions. This resulted in four systems: VLP_PL_-1 (∼2,000 lipids in total), VLP_PL_-2 (∼1,000 lipids in total), VLP_DG_-1 (∼2,000 lipids in total), VLP_DG_-2 (∼1,000 lipids in total) as shown in **Fig. 4C**. For each system, we performed 500 ns long equilibration simulation with the positions of protein backbone atoms restrained, which was followed by three independent 2,000-2,500 ns unrestrained simulations.

**Figure 4.**
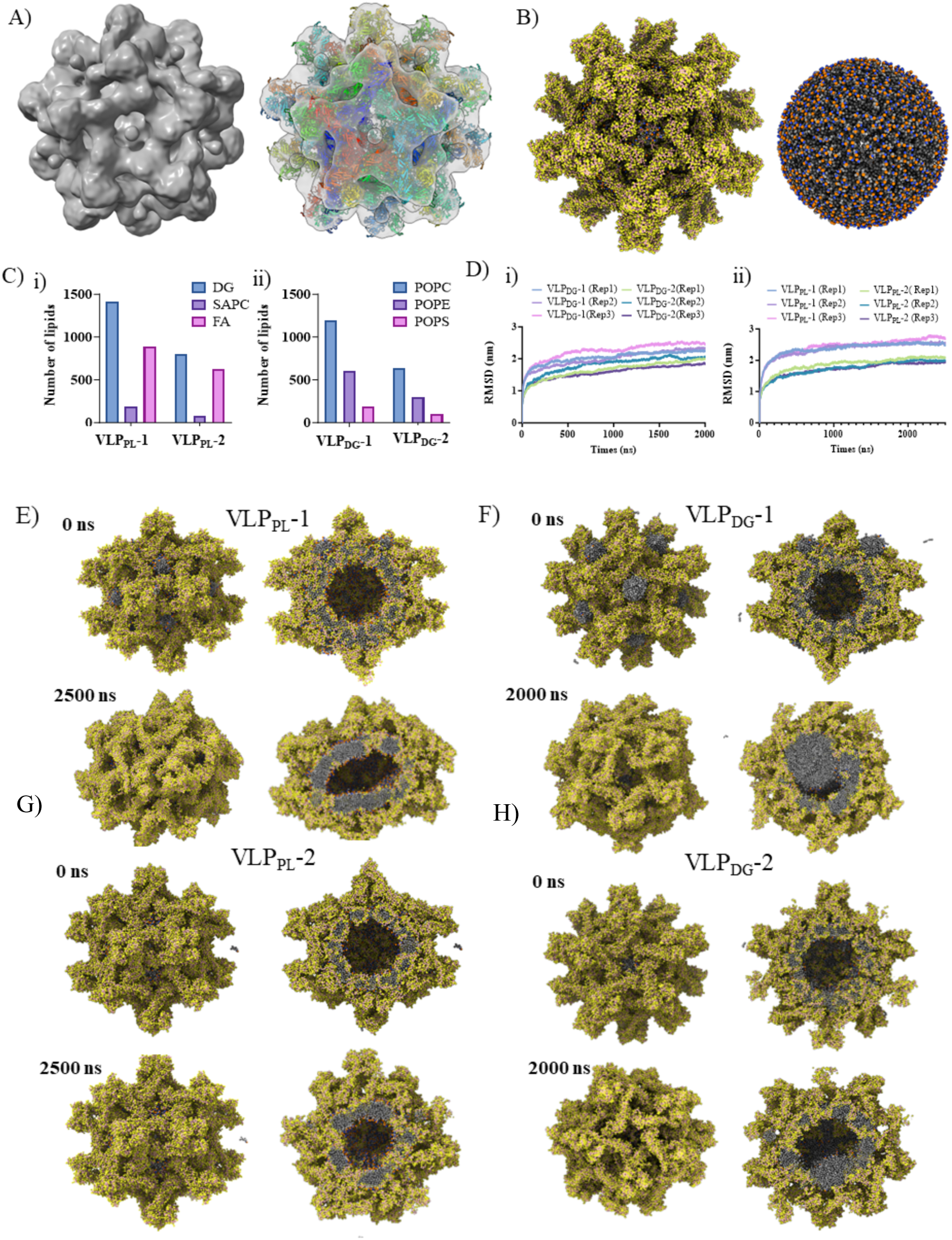
Coarse-grained MD simulations of imD2VLPs. **(A)** Cryo-EM map (at contour level=0.225) of imD2VLPs shown in surface representation (left) along with atomistic coordinates of 20 trimeric units of prM-E protein fitted into the cryo-EM density map (right). E-prM protein shell is shown in cartoon representation **(B)** Representative CG model of fully assembled imD2VLPs (left) together with lipid vesicle (right). **(C)** The lipid composition and number of lipid molecules in each VLP construct. **(D)** Root mean square deviation (RMSD) plot of VLP backbone beads with respect to the experimental structure is shown over the simulation time of VLP_DG_ and VLP_PL_ models respectively. **(E-H)** Initial and final snapshots of immature VLP simulations in PL and DG dominant lipid vesicles respectively. In **(B)**, **(E)**, **(F), (G),** and **(H)** the protein and lipids are shown as spheres (protein, lipid tails, and lipid headgroups colored in yellow, grey, and red, respectively).

The RMSD of all protein backbone beads for each VLP construct exhibited values ranging between ∼1.5-2.5 nm with respect to the experimental structure by the end of the simulation time (**Fig. 4D**). Among all the VLP simulation systems, VLP_PL_ constructs showed lower RMSD in comparison to VLP_DG_ systems. VLP constructs with a smaller number of lipid molecules (VLP_PL_-2 and VLP_DG_-2) tended to be more stable in comparison to systems with a higher number of lipids (VLP_PL_-1 and VLP_DG_-1), while the VLP_PL_-2 was the most stable amongst all systems (**Fig. 4E-H**). In the case of the VLP_DG_ simulation trajectories, formation of a lipid bulge in the lipid envelope was observed among all DG-containing VLP systems. The lipid bulge likely results from the previously observed accumulation of DG lipid molecules in the space between the lipid bilayer (lipid lens) arising from the trans-bilayer activity of DG-rich lipid vesicles^57,58^. Lipid lens formation was observed within the first 200 ns of the unrestrained production run (**Fig. 4H**). Such bulge formation among VLP_DG_ simulations was previously observed in both experimental and computational studies with mature VLPs (reference to be added, manuscript under review). The imD2VLP cryo-EM structure was obtained with icosahedral symmetry constraints and showed a spherical lipid bilayer (**Fig. 3 and 4**). In this case, we compared the more stable VLP_PL_ constructs obtained from our simulations. The final frames of VLP_PL_-1 and VLP_PL_-2 attained an ellipsoidal or spherical shape respectively, which indicated that the number of lipid molecules was a potential factor contributing to heterogeneity in the shapes observed in cryo-EM micrographs (**Fig. 4E and G**). Due to the low resolution of cryo-EM map, the stem-helix (SH) and TM regions in the imD2VLP structure were not visible and were modelled based on other experimental structural data (details in Methods). Analysis of the simulated dynamics of the stem helix (SH) and TM regions showed considerable differences between VLP_PL_-1 and VLP_PL_-2. The distribution plot of solvent accessible surface area (SASA) of each E protein monomer showed variations among SH and TM regions within each and between the respective VLP_PL_ MD simulations (**Fig. 5A; Supplementary Fig. 3**). The number of contacts between the lipid envelope and each SH region of E protein showed a significant difference between VLP_PL_-1 and VLP_PL_-2 due the different morphology that VLP constructs adopt during simulations (**Fig. 5B**). This suggests that each SH and TM region independently adopts a different orientation, and thus during EM-image reconstructions these regions would have been averaged out resulting in lack of clear density. PCA was performed on the concatenated triplicate simulation trajectory of each VLP_PL_ system and revealed the dominant motions exhibited in VLP_PL_-1 and VLP_PL_-2 trajectories. Comparing the most dominant motions (PCA1 and PCA2) which accounted for >70% of the covariance matrix, a higher magnitude of structural change in VLP_PL_-1 compared to VLP_PL_-2 and loss of spherical shape was observed (**Supplementary Fig. 4A-D**). We also calculated the distance between each pr molecule and DII domain of each E protein monomer, and we observed that in the case of the VLP_PL_-1 system (**Fig. 5C**), more pr molecules dissociated due to the loss of sphericity (**Supplementary Fig. 4E**). Collectively, our data suggests that ∼1,000 lipids and the PL-dominant environment provide more stable and robust imD2VLP particles.

**Figure 5.**
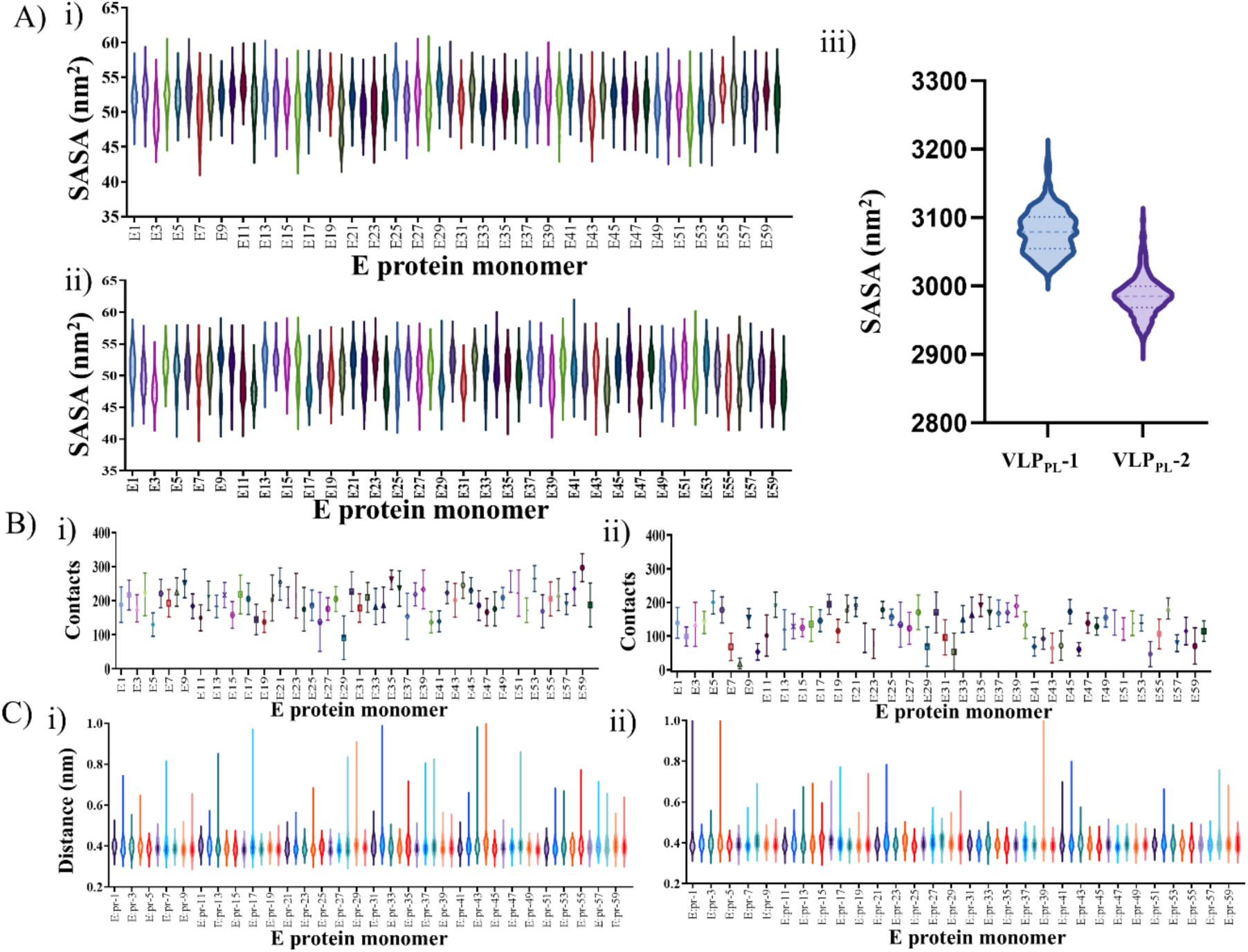
Dynamics of imD2VLPs in two lipid vesicle environments. **(A)** (i) Solvent accessible surface area (SASA) violin plots of per residue of stem helix (SH) region (residues 396 to 450) for each E protein monomer for VLP_PL_-1 (i) and VLP_PL_-2 (ii). (iii) SASA violin plot showing the distribution of all SH regions combined from VLP_PL_-1 and VLP_PL_-2. **(B)** Per-residue SH-lipids contacts for VLP_PL_-1 (i) or VLP_PL_-2 (ii) using 0.6 nm cutoff distance. (C) Violin plots showing the distributions of the minimum distance between DII and interacting pr peptides for each of 60 prM-E protein heterodimeric unit for VLP_PL_-1 (i) and VLP_PL_-2 (ii). All distributions and mean values were calculated over three simulation replicas.

### Targeted molecular dynamics (TMD) simulations capture the VLP maturation process

Similarly to the DENV virion, the imD2VLPs undergo large-scale conformational changes during maturation. The immature VLP transit from a spiky state with 20 trimeric protomers of prM-E protein to a smooth mature state with 30 units of dimeric E-M protein protomers. This is triggered by lowered pH and furin protease cleavage inside the host cell^59,60^. To explore the transition pathway during the VLP maturation process, we employed TMD simulations. In this approach, an additional bias is added to the total energy as a harmonic potential governed by the RMSD with respect to the target coordinates. The target RMSD evolves linearly over the simulation time and triggers the transition from the starting coordinates to the target structure^61^. Here, we employed TMD simulations of the VLP_PL_-1 as the initial immature trimeric state which undergoes maturation biased towards the smooth dimeric structure.

As described in the Methods section, steering forces were applied to specific E and M proteins to trigger the conformational transition. We superimposed the immature and mature VLP^20^ coordinates along the 5-fold symmetric axes and assigned each chain by the lowest RMSD value between the initial (immature) and final (mature) conformation (**Fig. 6A**). The overall RMSD value between the two states corresponds to ∼6 nm. Firstly, we assessed the influence of the strength of the RMSD bias applied to immature VLP beads. We tested four different force constants of the harmonic bias potential: 100, 500, 1,000, and 10,000 kcal mol-1 Å-2 (named TMD-1, TMD-2, TMD-3, and TMD-4, respectively) and ran four independent 100 ns long TMD simulations. In all TMD simulations, the conformational transition from trimeric spikey to dimeric smooth state occurred in a single run without any steric clashes between E protein monomers (**Fig. 6B**) while retaining a symmetrical particle shape. There were subtle differences observed across all TMD simulations depending on the force constant. The RMSD plot of all protein backbone beads over the simulation time with respect to the final mature state showed a non-linear change in RMSD in the case of TMD-1 when compared to the remaining systems. The TMD-1 simulation with lowest force constant showed a bimodal curve with respect to RMSD. Contrary to TMD-1, among other TMD simulations (TMD-2 to TMD-4), the gradual reduction in RMSD was linear over time (**Fig. 6Ci**). Also, we observed subtle changes in the lipid envelope during the transition among all the TMD simulations. The lipid vesicle diameter gradually changed over the simulation time across all the TMD simulations (**Fig. 6Cii**), from ∼250 Å to 242 Å in agreement with experimental values of 245 Å.

**Figure 6.**
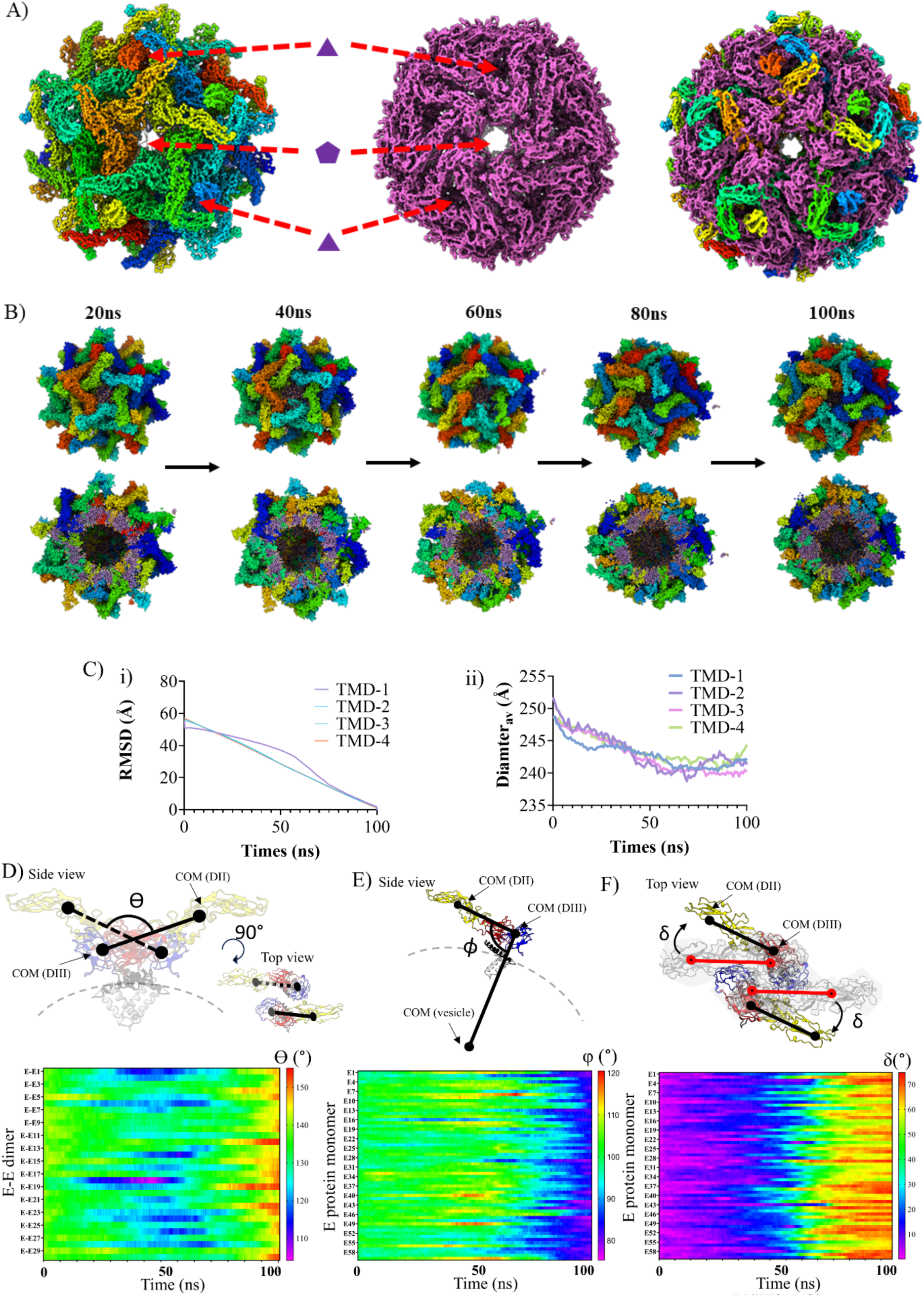
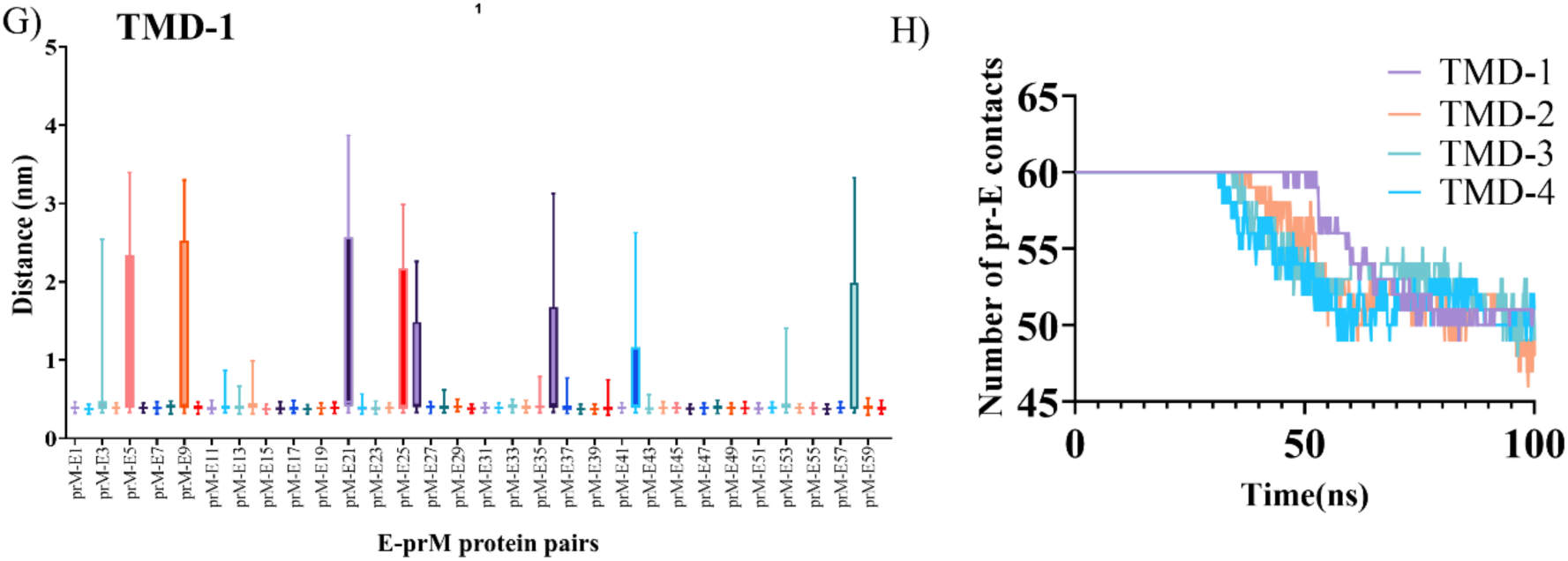
Targeted Molecular dynamics simulations capture mature to immature transition pathway of dengue VLP. **(A)** Immature (left, current study) and mature (middle, ^20^) VLP structures after fitting showing E protein alone with corresponding 5-fold and 3-fold symmetry axes. Each E protein monomer from immature and mature states are shown in surface representation and color-coded. Superimposed E protein only mature (purple) and immature (rainbow) VLP structures along the symmetry axes of VLPs (right). **(B)** TMD-1 simulation snapshots showing the transition from spiky trimeric state to smooth dimeric conformation. The entire VLP (top) and its cross-section (bottom) of immature VLP at t=20, 40, 60, 80 and 100 ns during the TMD simulation with protein and lipid beads shown in rainbow and purple CG-beads. **(C)** RMSD of protein backbone beads with respect to the final mature VLP structure over the simulation time for different force constants (100, 500, 1000 and 10000 kcal mol^-1^ Å^-2^) corresponding to TMD-1, TMD-2, TMD-3 and TMD-4 respectively (left). The VLP vesicle diameter over each TMD is shown on the right. **(D)** Cartoon representation of single unit of E protein dimer and angle ϴ along with the heatmap showing ϴ for all 30 pairs of E-dimers over the course of the TMD-1 simulation time (bottom). **(E)** Cartoon representation of E protein monomer and angle Φ along with the heatmap showing changes in angle Φ for all 60 E proteins during the TMD-1 simulation time (bottom). **(F)** Schematic showing single unit of E protein dimer at t=0 (grey shade) and later time point (colored) shown in cartoon representation from immature VLP. The reference line connecting DII and DIII at t=0 and same line at later time point shown in red and black respectively along with angle δ. The heatmap shows changes in angle δ for all 60 E-proteins during the TMD-1 simulation time (bottom). The E proteins are shown in cartoon representations and color coded in a domain-wise manner as described previously with stem in black and TM in white **(D)-(F)**. **(G)** Box plot showing the minimum distance between domain II and pr peptides of each E-prM protein heterodimer from TMD-1 simulation trajectory. **(H)** The number of pr peptides within 0.6 nm cutoff distance of respective monomer DII.

Overall, all the TMD simulations showed a smooth transition of trimeric to dimeric state via a sliding-rotating motion without any steric clashes among E-prM protein (**Supplementary Movie 2**). This agrees with a recently proposed maturation model for DENV described by Duquerroy et. al using icosahedral crystal lattices at low pH and neutral pH^62^. To quantify this sliding and rotating motion, we measured three structural features i.e., angles either within each E protein monomer or between dimers over the course of the simulation time (see Methods section). All three angles measured in the first frame correspond to the experimentally determined cryo-EM structure and subsequent changes reported relate to the structural changes occurred during the TMD simulations (**Supplementary Table 3**). Angle ϴ measured and averaged over the 30 pairs of E-dimers in the first frame of the simulation shows 142°±0.2° and attains a final value between 148°±0.3°-160°±0.1° in the final frame of the simulation (mature state) depending on force constant used in given TMD simulation (**Supplementary Table 3**). Interestingly, for almost all the E protein pairs and from all four TMD simulations, the ϴ value gradually drops to ∼120° from the initial value and starts to increase in the latter part of the simulation time (**Fig. 6D**). This shows that during the sliding-rotating motion, the E-dimers perform a “scissor”-like motion during the transition from spiky immature to a spherical mature state. In the case of Φ, the angle gradually decreased from 98°±0.2° to 82°±2° in TMD-1 or to 74°±0.6° in the case of TMD-4 (**Supplementary Table 3**). It is noteworthy that the averaged ϴ and Φ values for the whole imD2VLP from the final frame of TMD simulations show comparable values with respect to that of the mature VLP. In particular, the final state of TMD-4 simulations attains an identical ϴ (∼160°) value averaged over all the 30 dimeric units of mature VLP structure **(Supplementary Table 3)**. Finally, angle δ over the simulation time showed that each E protein monomer gradually adopted an angle of 65°±5° to 68°±4° with respect to the initial E protein orientation at t=0 in the immature state. Further, we measured the distance between pr domain of prM protein and DII of E protein among all E-prM protein pairs. This showed that most of the prM-DII pairs stay in contact except for a few events of displacement during the transition (**Fig. 6G-H; Supplementary Fig. 5**). The number of pr domain molecules dissociating from E proteins remained similar (∼10) irrespective of the force constant used and agrees with unbiased MD simulations (**Fig. 5C; Supplementary Fig. 4E**). This suggests that during TMD simulations, the “pulling” rate was not playing a major role in pr-E protein interactions in VLP system. In the cellular context, previous studies on DENV have shown that exposure to low pH (∼6) in the Golgi-network drives the structural changes resulting translation of E-prM protomers to form flat dimers from spiky trimers, covering the whole lipid envelope encasing the nucleocapsid. In the case of VLPs, our TMD simulations showed that translation i.e., sliding of each prM-E protomers accompanied with rotation, forms quaternary contacts to assemble into a smaller icosahedral particle.

### Empty cryo-EM density maps linked to dynamic lipids and N-glycans

During the equilibration simulations of both VLP_PL_ and VLP_DG_ systems, we observed that some lipid molecules spontaneously protrude out of the spherical lipid vesicle (**Fig. 7A**) at the 5-fold symmetry axes in all simulations irrespective of lipid composition and number of lipids used in the vesicle. Further, these lipid molecule protrusions reintegrated into the lipid vesicle within 10 ns of unrestrained production simulations. Nonetheless, 1-2 lipid molecules were observed to escape the VLP assembly into the bulk solvent during the unrestrained simulations across all replicates. Upon closer inspection of the cryo-EM density map, we consistently observed a low-resolution density emerging out of the lipid vesicle density at all the 5-fold axes (**Fig. 7B**). This extra non-protein density was likely due to protruding lipid molecules captured by the EM reconstruction due to particle averaging. This indicates that the protrusion and escape of lipid molecules occurring in solution with immature VLPs was captured by cryo-EM particle averaging during image reconstruction and MD simulations.

**Figure 7.**
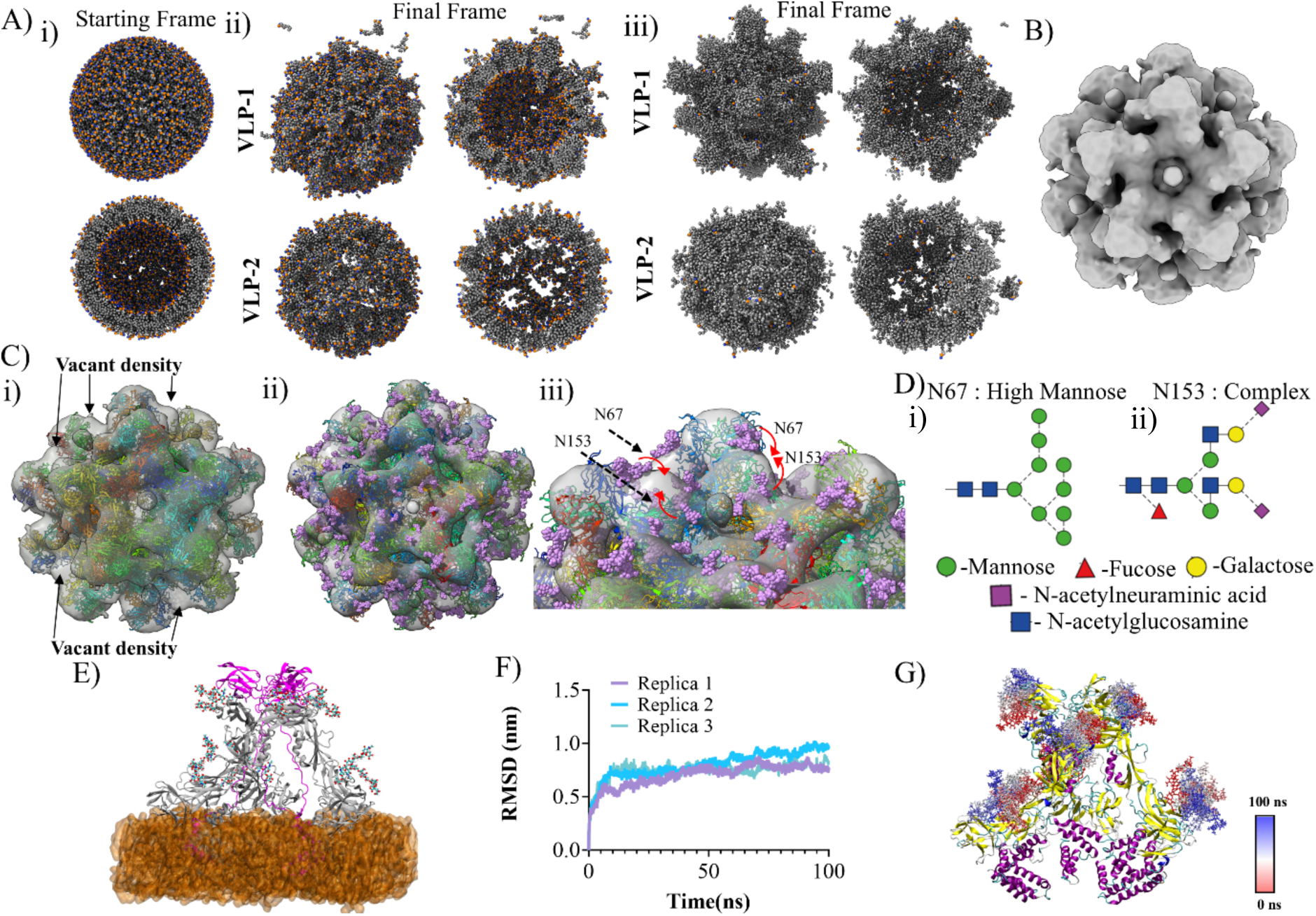
MD simulations link the unoccupied regions post prM-E protein shell fitting to the cryo-EM density map of imD2VLP fitting to dynamics of glycans and lipids. **(A)** (i) Representative CG PL dominant lipid model visualized together with its cross-section. ii-iii) Snapshots of lipid envelope from first and last frame of 500ns long equilibrium simulation of VLP_PL_-1 (ii) and VLP_PL_-2 (iii) shown in whole and in cross-section. **(B)** Cryo-EM map of immature DENV shown in surface representation with magnitude of contour at 0.27**. (C)** (i) Glycan-free and (ii) glycosylated atomistic model of immature DENV VLP fitted into the cryo-EM density map with unoccupied cryo-EM density regions highlighted. (iii) “Zoom in” on the two glycosylated trimeric spikes. **(D)** Glycan composition and connectivity at N67 and N153. **(E)** Initial coordinates of the trimeric glycosylated E-prM protein embedded in a DG -dominant lipid bilayer. Each protein and lipid molecule shown in cartoon and surface representation respectively (E proteins: grey; prM protein: magenta and lipid bilayer: orange). **(F)** RMSD of the protein backbone atoms over the course of three independent MD simulation replicas. **(G)** Starting structure (t=0) of trimeric E-prM protein along with the simulation snapshots of glycans at N-67 and N-153 with their coordinates shown at every 10ns over the course of 100ns simulations. Each protein molecule represented in cartoon representation and color coded as per secondary structure (yellow: β-sheets, magenta: α-helices, cyan: loops or unstructured regions). Glycans are shown in stick representation and color coded as per simulation time t=0 (blue) to t=100 (red).

Another region of unoccupied cryo-EM density was observed after fitting the atomistic model of the VLP into the experimental density map (**Fig. 7C**). We observed that this region of the map was juxtaposed to the N67 and N153 sites of E protein trimers (**Fig. 7Di-ii**). We modeled experimentally derived glycans^45^ on each protein trimer and fitted these coordinates into all-atom VLP coordinates (**Fig. 7E**). The modeled glycans occupied the empty space of the cryo-EM map (**Fig. 7Di**). The all-atom MD simulations of glycosylated trimeric E-prM protein embedded in the lipid bilayer containing DG:PC:FA (56:28:6) revealed highly dynamic glycans at both N67 and N153 sites with a large average volume occupied (**Fig. 7F-G; Supplementary Movie 3**). An average density map of imD2VLPs generated by combining the structural fluctuation of unrestrained MD simulations for the VLP_PL_ systems of whole immature imD2VLPs did not show any extra density as our VLP models were not glycosylated (**Supplementary Fig. 4C-4D**). Therefore, this likely explains the unoccupied areas observed in the cryo-EM map.

## DISCUSSION

Our previous work elucidated and characterized the structure of mature dengue serotype 2 virus-like particles (mD2VLPs) and their potential to elicit a robust immune response^20^. This demonstrated that these particles possess antigenic structures conducive to revealing cryptic epitopes, thereby enhancing immunogenicity and offering protection against all four dengue serotypes in murine models. Building on this foundational work, we have now elucidated the structure of immature DENV VLP to comprehensively understand the global conformational changes involved in the maturation process as well as the native intrinsic dynamics of VLPs governing the stability.

One central question was how low pH in the trans-Golgi induces the necessary conformational rearrangements in the spike of the VLP system. Although the 9 Å resolution of the cryo-EM map limited atomic-level interpretation, the integration of cryo-EM data with CG simulations provided critical insights into the dynamic nature of VLP maturation. These simulations revealed a sliding-rotating motion of E protein dimers during the transition from an immature trimeric state to a mature dimeric state **(Fig. 8; Supplementary Movie 2)**, offering molecular-level detail into the structural choreography of maturation. This detailed understanding is crucial for the rational design of VLPs, particularly as immature particles have a higher propensity to generate ADE-causing antibodies, which is a major concern for vaccine safety.

**Figure 8.**
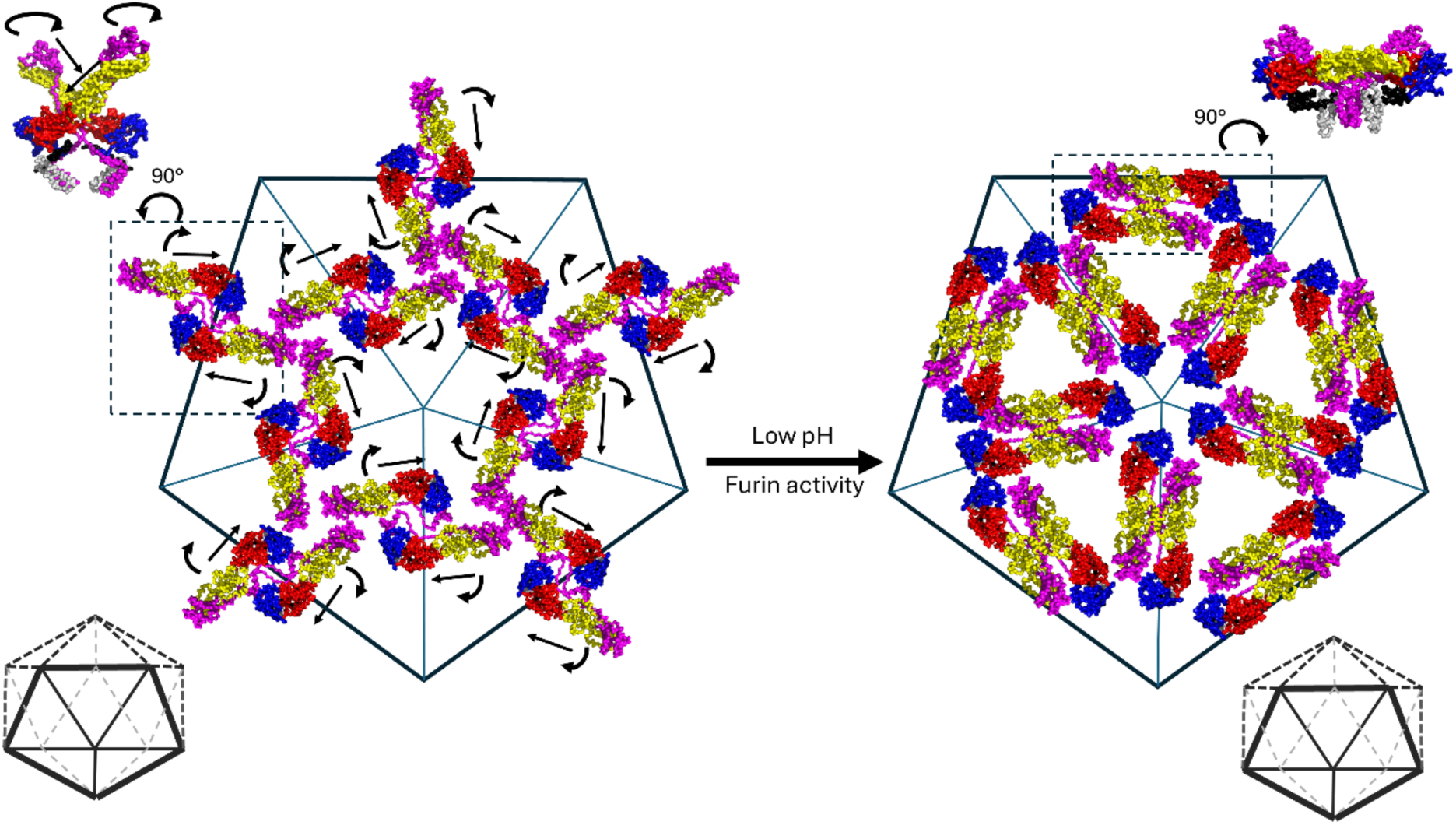
Schematic illustration of structural transition of E-prM protein unit from immature to mature state. Each particle consisting of 60 units of E-prM protein arranged in T=1 icosahedral symmetry with 20 units at 3-fold symmetry axes and 12 units at 5-fold symmetry axes. E-prM protein units at 5-fold axis highlighted undergoes “sliding –rotating” (pink arrows) motion as shown in the TMD simulation from spiky immature state to flat spherical mature state. Inset shows the side view of dimeric pr-M protein unit in immature and mature particle. The E-prM proteins are color coded in a domain-wise manner (DI-red, DII-yellow, DIII-blue, stem-black, TM-white and prM-magenta) and shown in cartoon representation.

There have been several elegant structural studies proposing diverse maturation models for flaviviruses. The most recent cryo-EM study of dengue VLP maturation suggested transition which involves substantial structural changes ^63^. Another study on DENV by cryo-EM structure suggests that the prM linker acts as a “drawstring,” pulling on the ectodomains to rotate them^24^. Similarly, the cryo-EM structure of immature Spondweni virus (SPOV) posits that histidine protonation on the prM linker allows the E protein to transition from a strained, spiky state to a smooth conformation via collapsing mechanisms^64^. Alternatively, an immature virus cryo-EM structure of Binjari virus (BinJV) indicates the full structure of the prM linker at 4.4 Å resolution^54^ and proposed the “pillar-collapse” model. Moreover, a recent cryo-EM study on immature ZIKV proposes the “latch and lock” mechanism for flavivirus maturation. This mechanism, driven by electrostatic and hydrophobic interactions between M and E proteins, orchestrates the structural rearrangements essential for maturation, ultimately stabilizing the virion for subsequent infection^65^. These models highlight the complexity and diversity of maturation mechanisms across flaviviruses.

Interestingly, the basic protomer of both the immature dengue virion and immature dengue VLPs remain the same i.e., spiky trimeric E-prM protein. In the mature state, the virion and mature dengue VLPs are the E-M protein raft consisting of trimer of E-M protein dimers and E-M protein dimers respectively. It is therefore highly plausible that the E-prM spiky assemblies from immature dengue virion undergo sliding-rotating motion to attain flat dimeric state akin to immature dengue VLPs. This suggests that the “pulling” mechanism may not play a major role in pr-E protein interactions during the maturation process. However, given the higher number of E-M protein units (180 copies) in virion compared to VLP (60 copies), we cannot rule out the possibility that the maturation dynamics of the virion involve more complex interactions and conformational states^54,64,66^.

A key question pertains to the immunogenic potential of flaviviral VLPs compared to virions. Our dengue VLP cryo-EM analysis unveiled a complex architecture, characterized by a glycoprotein layer adorned with 20 prominent spikes consisting of trimeric E-prM protein embedded in lipid envelope. These spikes exhibited a T=1 arrangement and were composed of three pairs of E-prM heterodimers capped by three globular pr domains. Noteworthy similarities to natural immature flaviviruses underpins the authenticity of our imD2VLPs (**Supplementary Fig. 6**). The observed steric hindrance at domain III (DIII) and around amino acid 101 in the fusion loop suggests potential differences in antibody accessibility between imD2VLPs and mD2VLPs, with important implications for immune response modulation. By understanding these structural features, we can fine-tune VLP design to reduce the risk of ADE and improve vaccine efficacy.

Further, CG MD simulations allowed us to probe the functional dynamics of imD2VLPs, offering insights into their stability and behavior with respect to lipid envelope composition and total number of molecules. Our results indicated that imD2VLP stability was influenced by both lipid compositions where fewer lipids and PL dominant vesicles retained more spherical and robust morphology. Conversely, the formation of a lipid bulge in the case of DG dominant lipid envelopes raises questions about the stability and integrity of imD2VLP membranes. These observations are critical for improving VLP production methods, where lipid composition must be carefully controlled to maintain particle integrity.

Additionally, the dynamic behavior of lipid molecules observed in both CG and all-atom MD simulations could be linked to empty densities observed in cryo-EM maps. Lipid protrusions observed at the five-fold symmetry axes during MD simulations, and their correlation with the cryo-EM density, provide crucial insights into lipid dynamics. Understanding this behavior is essential for further refining the design of VLPs, particularly in terms of membrane stability and antigen presentation.

While this study represents a significant advance in the understanding of dengue VLP maturation, several limitations must be acknowledged. The resolution of the cryo-EM structure limited our ability to resolve fine molecular details. Addressing particle heterogeneity and improving structural resolution through techniques such as cryo-electron tomography will be essential in future studies. Additionally, the variation in VLP conformations and properties due to differences in expression and secretion systems warrants further investigation. Finally, in vivo studies evaluating the immunogenicity and protective efficacy of optimized VLP designs will be crucial for advancing vaccine development efforts.

Collectively, our study enhances the understanding of dengue VLP maturation and demonstrates improved VLP production efficiency in a mammalian cell system, aiding in the development of strategies to enhance the maturation process for better immunogenic properties. The insights into the structural dynamics of imD2VLPs provide a foundational understanding of surface protein rearrangement, enhancing our capacity to predict and potentially manipulate the immunogenic properties of dengue VLPs.

## Availability of data and materials

The EM structures have been deposited in the Electron Microscopy Data Bank (EMDB ID code:XXXX).

## Acknowledgements

The cryo-EM experiments were performed at the Academia Sinica Cryo-EM Facility (ASCEM) in Taiwan and International Institute for Macromolecular Analysis and Nanomedicine Innovation (i-MANI) at National Cheng Kung University in Taiwan. The computational work for this article was fully performed on resources of the National Supercomputing Centre, Singapore (https://www.nscc.sg).

## Authors contributions

S.-R.W., D.-Y.C. and J.-K. M. conceived the project. S.-R.W., D.-Y.C., J.-K. M., G.-W.C., Y.-N. W. conducted the experiments. G.-W.C., C.-H.W. contributed to the cryo-EM data acquisition and analysis, and S.-R.W. performed the three-dimensional reconstruction of the images. S.-R.W., D.-Y.C., J.-K. M., G.-W.C., Y.-N. W., V.-R. P., P.-J. B. interpreted and discussed the results, forming the theoretical framework. S.-R.W., V.-R. P. and J.-K. M. compiled the data and wrote the manuscript. All authors contributed to the revision of the article and approved the submitted version.

