## Supplementary Materials for "Structural Insights into Immature Dengue Virus-Like Particles Revealed by Cryo-EM and Molecular Dynamics Simulations"

### Legends

**Movie1:** Movie showing the 360° view of atomistic model fit to the Cryo-EM density of immature VLP particle.

**Movie2:** Trajectory showing the immature to mature transition of VLP during the targeted molecular dynamics simulation (TMD-1).

**Movie3:** 100 ns all atom simulation of E-prM trimeric protein embedded in DG dominant lipid bilayer. Protein is shown in cartoon representation along with glycans at N67 and N153 shows stick-sphere .Membrane lipids are shown as black spheres.

**Supplementary Table 1. Immature dengue serotype 2 virus-like particles (imD2VLPs) CG model simulation setup**

| E-protein mutation | Box Size (nm) | Total number of lipids | Lipid membrane composition | NaCl concentration | Production run (ns) |
|---|---|---|---|---|---|
| $VLP_{PL}$-1 | 41.5×41.5×41.5 | ~2000 | POPC:POPE:POPS (60:30:10) | 100mM | 3 × 2500 ns |
| $VLP_{PL}$-2 | | ~1000 | | | |
| $VLP_{DG}$-1 | 41.5×41.5×41.5 | ~2000 | DG:SAPC:FA (56:26:8) | | 3 × 2000 ns |
| $VLP_{DG}$-2 | | ~1000 | | | |

**Supplementary Table 2. All atom glycosylated E-prM prM protein simulation setup**

| Box Size (nm) | No. of water molecules | Lipid membrane composition | NaCl concentration | Production run (ns) |
|---|---|---|---|---|
| 15.5×15.5×18.3 | 100,536 | DG:SAPC:FA (56:26:8) | 100mM | 3 × 100 |

**Supplementary Table 3. Comparison of φ, δ and Ɵ angles between starting and final frames of all TMD simulations. The values shown are averaged over the entire simulation time (av) error bars correspond to standard deviations**

| System | Simulation | Force constant ( kcal mol$^{-1}$ Å$^{-2}$) | RMSD (Å) at final frame (t=100ns) | $\Phi_{av}$ (t=0) | $\Phi_{av}$ (t=100ns) | $\Theta_{av}$ (t=0) | $\Theta_{av}$ (t=100ns) | $\delta_{av}$ (t=0) | $\delta_{av}$ (t=100ns) |
|---|---|---|---|---|---|---|---|---|---|
| **Immature VLP-1** | **TMD-1** | 100 | 11.0 | 98.28±0.17 | 82.08±2.3 | 142.3±0.23 | 147.7±3.1 | 1.61±0.7 | 64.6±5.0 |
| | **TMD-2** | 500 | 5.7 | | 76.22±1.4 | | 156.6±1.8 | | 67.0±5.9 |
| | **TMD-3** | 1000 | 4.9 | | 75.09±0.9 | | 158.3±1.1 | | 67.8±7.3 |
| | **TMD-4** | 10000 | 2.8 | | 74.30±0.6 | | 160.1±0.8 | | 68.3±4.2 |
| **Mature VLP-1** | **-** | - | - | - | 106.0±0.7 | - | 160.9±0.9 | - | - |

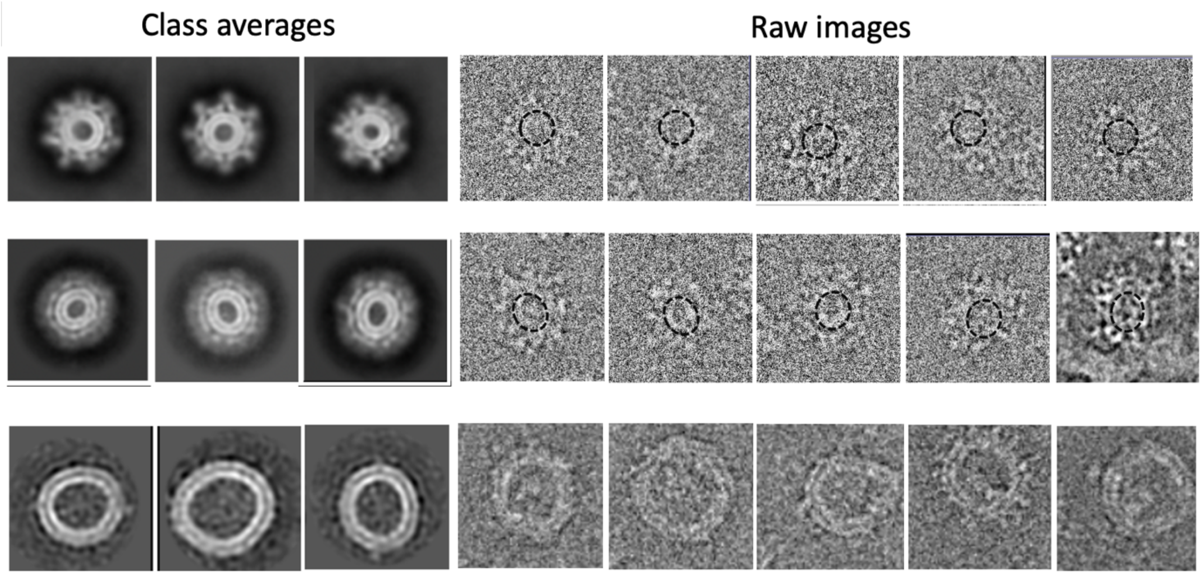


**Supplementary Figure 1. Representative 2D class averages from reference-free classification.** There are more raw images listed. The left panel lists the class averages while the right panel lists the raw images.

`

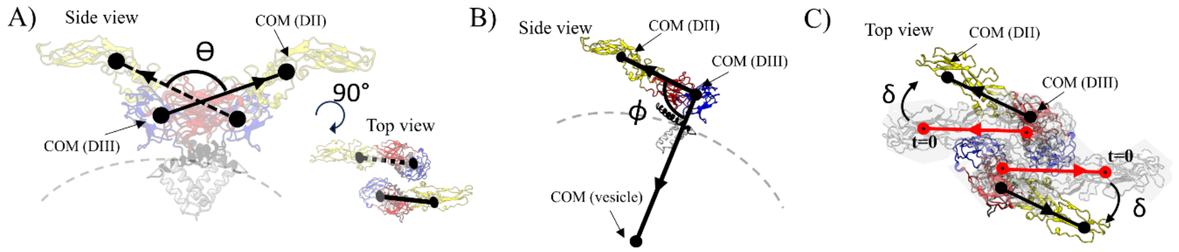


**Supplementary Figure 2. (A)** Cartoon representation of single unit of E protein dimer from immature VLP along with vectors connecting COMs of DII and DIII forming angle Θ are shown. **(B)** Cartoon representation of E protein monomer from immature VLP along with vector connecting COMs of DII-DIII, DIII-lipid vesicle and angle Φ are shown. **(C)** Schematic showing single unit of E protein dimer at t=0 (grey shade) and later time points (colored) shown in cartoon representation from immature VLP. The reference vector connecting DII and DIII at t=0 and later timepoint is shown in red and black respectively along with angle δ. The E proteins are color coded in a domain-wise manner (DI-red, DII-yellow, DIII-blue, stem-black, TM-white) and shown in cartoon representations.

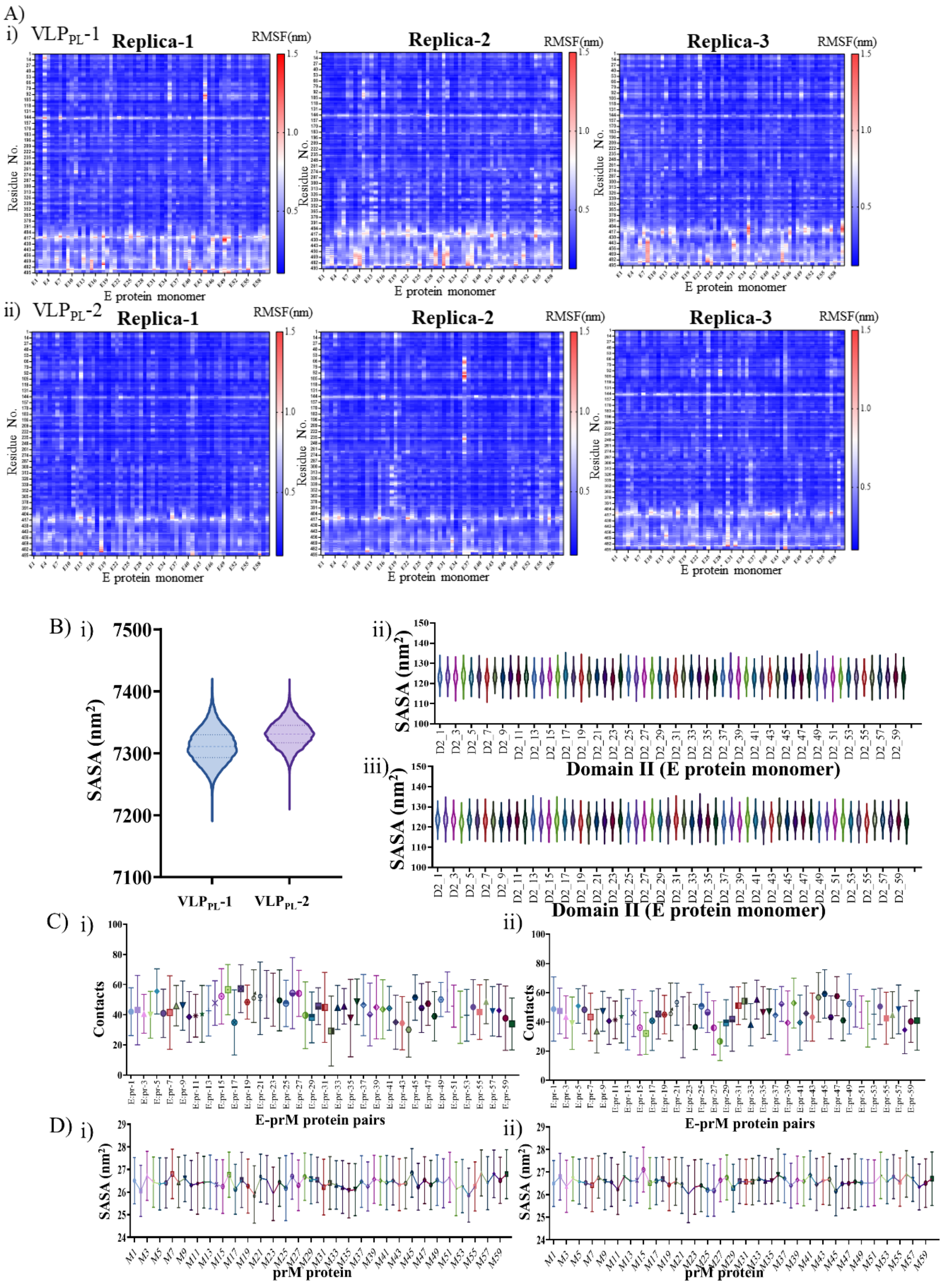
A)
i) VLP_PL-1
Replica-1
Replica-2
Replica-3
RMSF(nm)
Residue No.
E protein monomer
ii) VLP_PL-2
B) i)
SASA (nm²)
VLP_PL-1
VLP_PL-2
ii)
iii)
Domain II (E protein monomer)
C) i)
ii)
Contacts
E-prM protein pairs
D) i)
ii)
prM protein

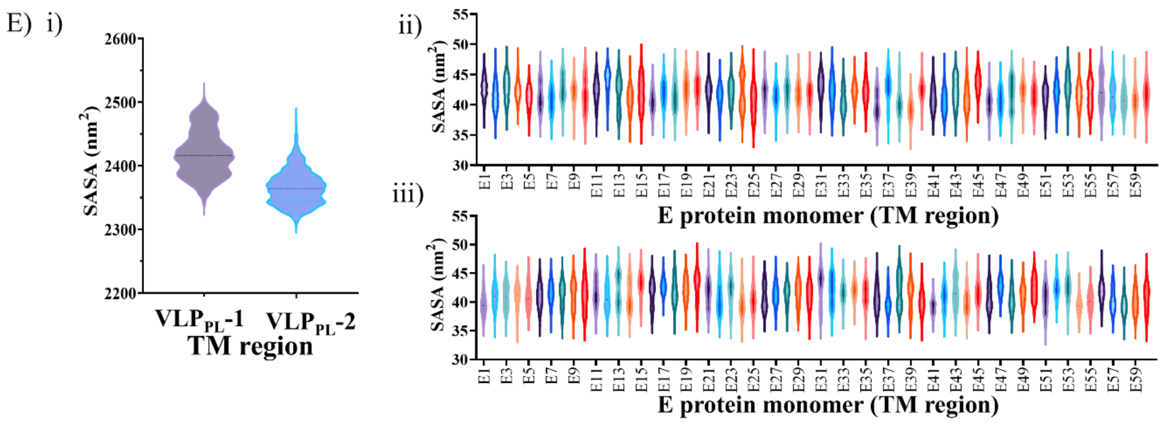


**Supplementary Figure 3. Comprehensive analysis of structural dynamics and interactions in VLP$_{PL}$-1 and VLP$_{PL}$-2 simulations. (A)** Heatmap showing per-residue RMSF values of each E protein from VLP$_{PL}$-1 and VLP$_{PL}$-2 simulation trajectories from three replicas. **(B)** Violin plot showing the distribution of SASA of DII from VLP$_{PL}$-1 and VLP$_{PL}$-2 from concatenated triplicate simulation trajectory of each 2500ns. (i) Plot showing the distribution of SASA for DII domain of each E protein monomer from VLP$_{PL}$-1 (ii) and VL$_{PL}$-2 (iii) from concatenated triplicate simulation trajectory of each 2500ns. **(C)** Average number of contacts between each pr domain: DII pair, plot shows mean and standard deviation values calculated after combining three independent 2500ns long simulations from VLP$_{PL}$-1 and VLP$_{PL}$-2 systems. **(D)** Plot showing average SASA value for each pr domain from VLP$_{PL}$-1 (ii) and VL$_{PL}$-2 (iii) from concatenated triplicate simulation trajectory of each 2500ns. **(E)** Violin plot showing the distribution of SASA of TM from VLP$_{PL}$-1 and VLP$_{PL}$-2.

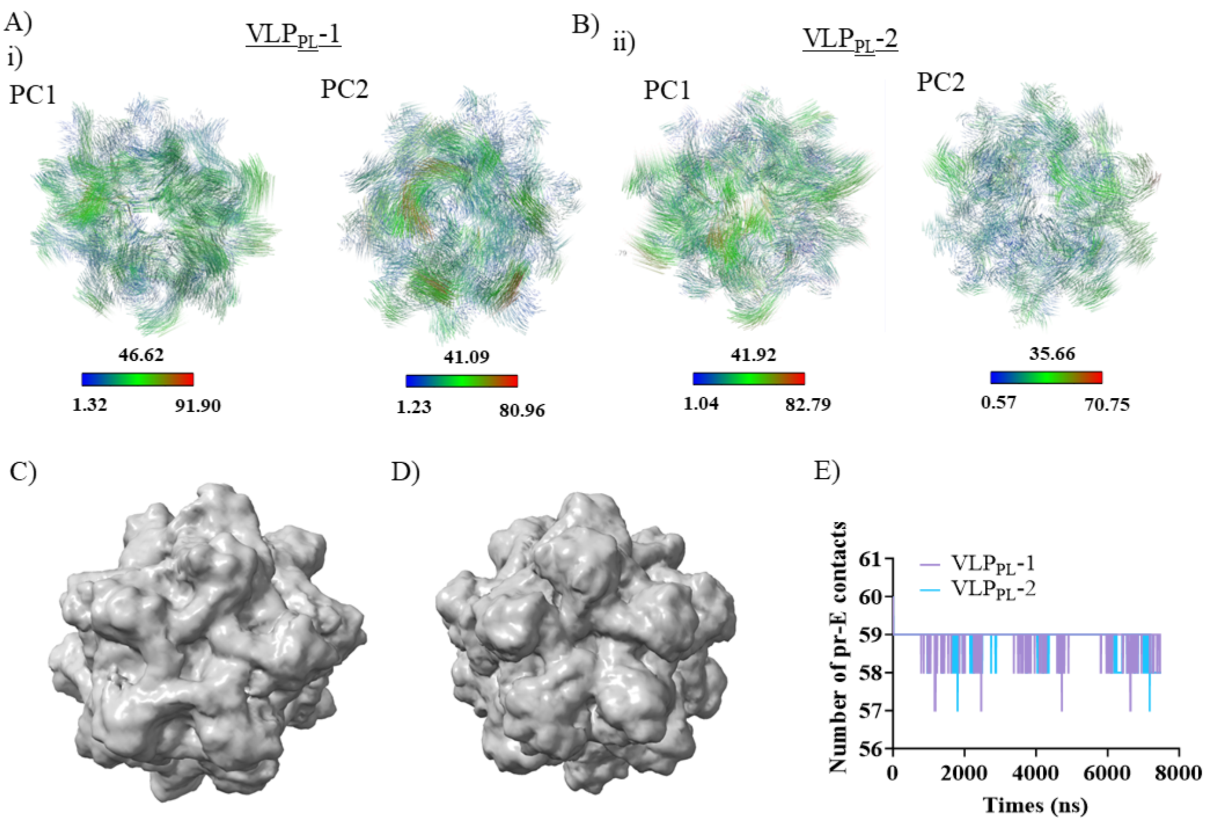


**Supplementary Figure 4**. **Analysis of principal component modes, structural fluctuations, and interactions in VLP$_{PL}$-1 and VLP$_{PL}$-2 simulations. (A-B)** First principal mode (PC1) motions from POPC dominant immature VLPs extracted from 7.5µs long simulation trajectory accounting for 45.02 and 41.4 % of variation respectively. **(B)** Second principal component mode (PC2) capturing the coordinated motions of each E-prM protein subunit. PCA was performed on every third protein back bone bead of each VLP trajectory and PC1 accounts for of ~45-50% variation in the data. **(C-D)** Surface representation of density generated from ensemble average of structural fluctuations calculated from concatenated triplicate VLP$_{PL}$-1 **(C)** and VLP$_{PL}$-2 **(D)** simulation trajectories. VMD volmap tool was used to generate the densities at a resolution of 5Å considering VLP structures at every 1ns from concatenative triplicate simulations. **(E)** plot showing number pr domains in contact with DII domain from 60 prM-E protein heterodimers in fully assembled VLP from VLP$_{PL}$-1 and VLP$_{PL}$-2 simulation of each 2500ns.

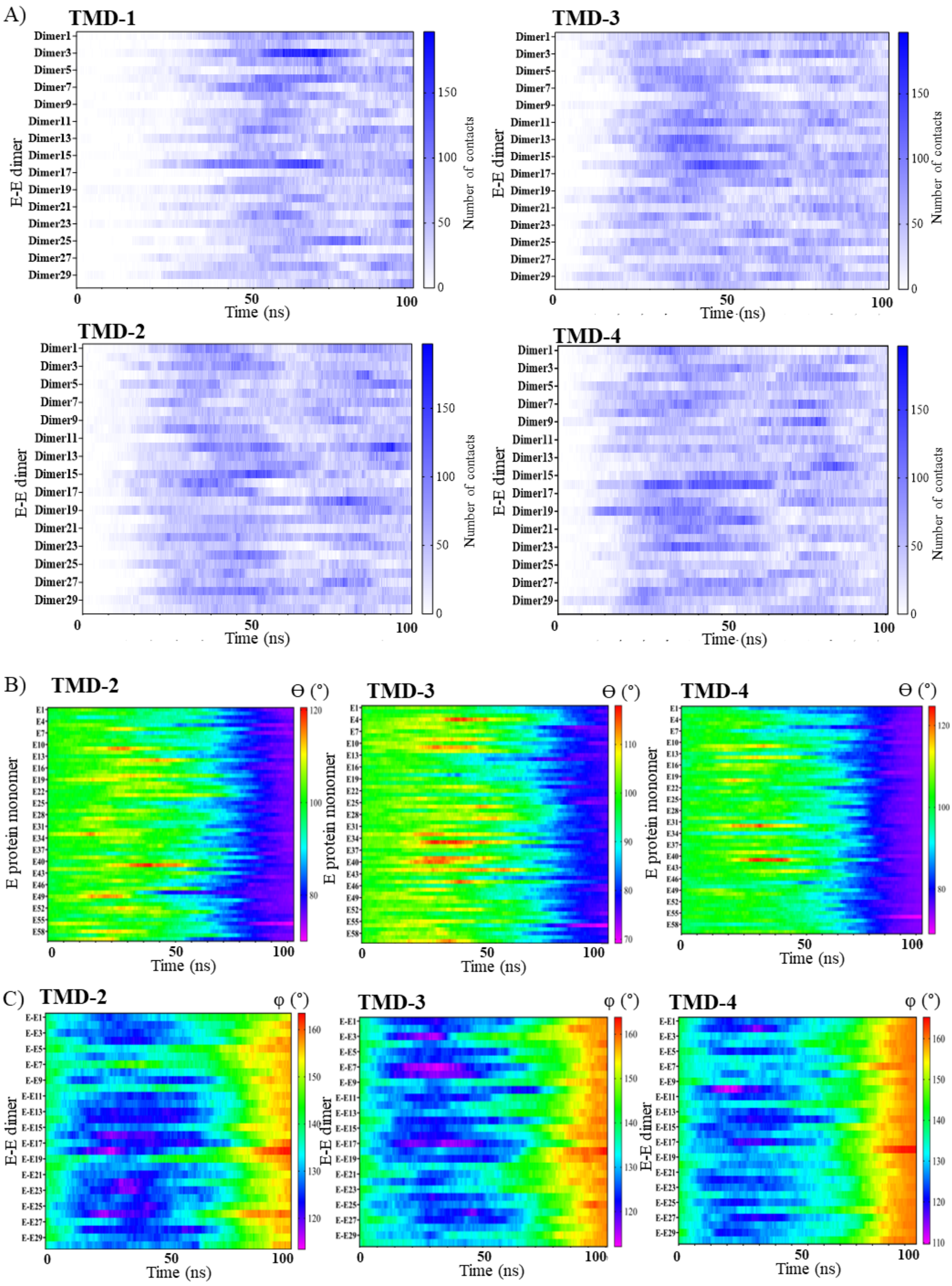
A)
TMD-1
TMD-3
TMD-2
TMD-4
E-E dimer
Dimer1
Dimer3
Dimer5
Dimer7
Dimer9
Dimer11
Dimer13
Dimer15
Dimer17
Dimer19
Dimer21
Dimer23
Dimer25
Dimer27
Dimer29
Number of contacts
0
50
100
150
Time (ns)
B)
TMD-2
TMD-3
TMD-4
Θ (°)
E protein monomer
E1
E4
E7
E10
E13
E16
E19
E22
E25
E28
E31
E34
E37
E40
E43
E46
E49
E52
E55
E58
Time (ns)
C)
TMD-2
TMD-3
TMD-4
φ (°)
E-E dimer
E-E1
E-E3
E-E5
E-E7
E-E9
E-E11
E-E13
E-E15
E-E17
E-E19
E-E21
E-E23
E-E25
E-E27
E-E29
Time (ns)

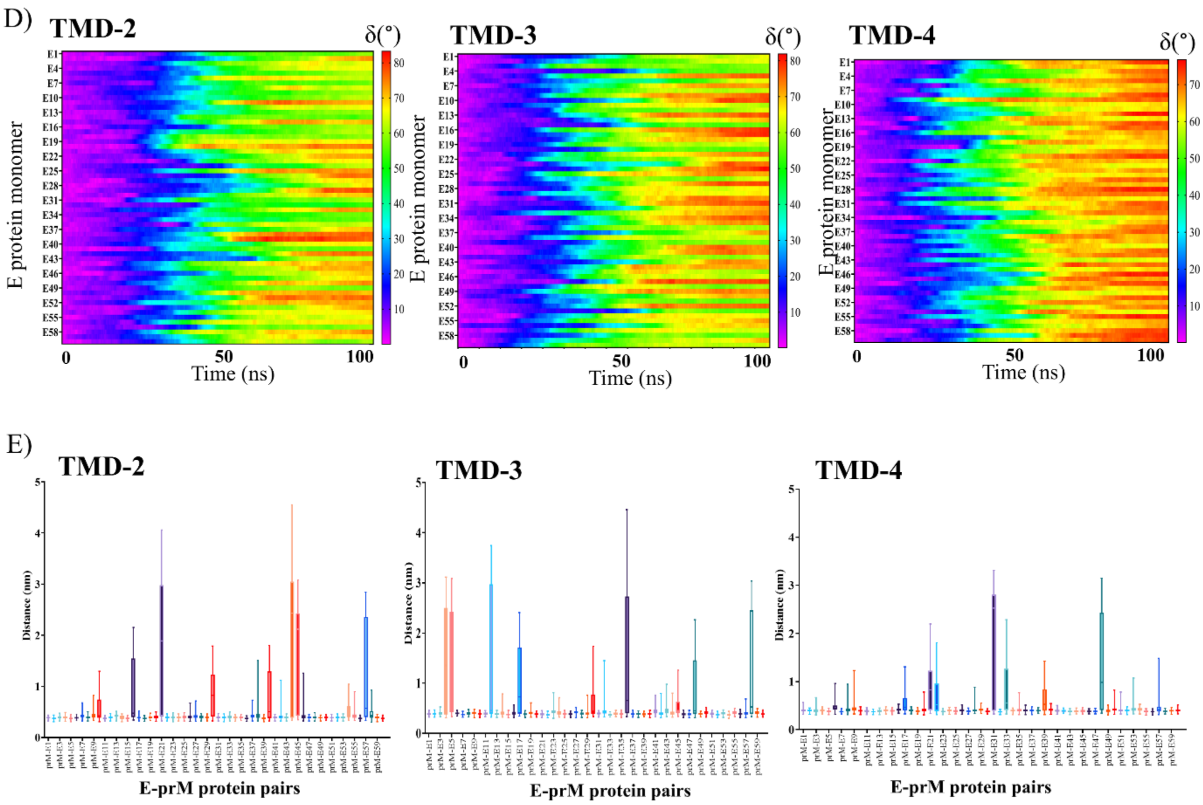


**Supplementary Figure 5. TMD simulations analysis. (A)** Heatmap showing the number of contacts between each E protein dimer with respect to time from all TMD simulations. **(B)** Heatmap shows the changes in angle ϴ between E protein dimers during the 100ns TMD trajectory for each E protein in TMD simulations. **(C)** Heatmap shows the changes in angle φ during the 100ns TMD trajectory for each E protein in TMD simulations. **(D)** Heatmap shows the changes in angle δ during the 100ns TMD trajectory for each E protein in TMD-simulations. **(E)** Box plot showing the distribution of distance between domain II and pr domain of E protein and prM protein respectively from all TMD simulations.

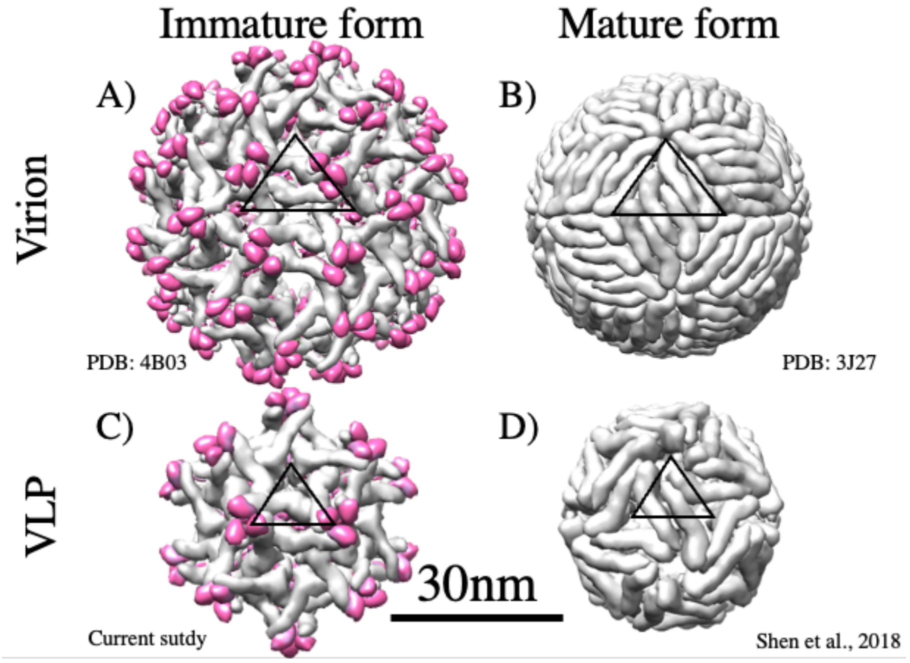


**Supplementary Figure 6. Structural arrangement of E and (pr)M proteins in different viral and VLP states.** Organization of the E and (pr)M proteins on the surface of **(A)** immature, **(B)** mature virus, **(C)** immature VLPs and **(D)** mature VLPs. E monomers are shown as a Cα backbone with DI, DII and DIII in red, yellow and blue, respectively. The black triangle represented an icosahedral asymmetric unit.
